# Novel Regulators of Sugar-dependent Food Consumption in *Drosophila melanogaster*

**DOI:** 10.64898/2026.11.02.763739

**Authors:** Mubaraq Opoola, Lucas Fitzgerald, Shakil Ahmed, Makayla Wright, Sarayu Alli, Nicholas Wright, Dae-Sung Hwangbo

## Abstract

Diet and genetic background interact to influence food consumption, yet the genetic basis of diet-dependent feeding behavior remains poorly understood. Here, we used genome-wide association analysis (GWAS) to investigate the genetic regulation of sugar-dependent food consumption in *Drosophila melanogaster*. Food intake was quantified in 176 lines of the *Drosophila* Genetic Reference Panel (DGRP) using the Con-Ex (<u>Con</u>sumption-<u>Ex</u>cretion) assay on low- and high-sugar foods. Flies showed compensatory feeding, consuming more low-sugar food than high-sugar food, and this behavior was largely maintained across genotypes. Additionally, the magnitude and overall consumption varied several-fold among lines, with broad-sense heritability of 0.53 to 0.66 across sexes and diets. Association analysis across six sex- and diet-specific phenotypes identified 161 variants mapping to 118 candidate genes, which shared almost no loci between dietary conditions, indicating that consumption on the two foods is largely under separate genetic control. RNAi knockdown of candidate genes identified *CG42673* and *Tmtc2*, whose knockdown increased consumption across sexes and diets and was reproduced in independent experiments. Together, these findings show that natural genetic variation contributes substantially to diet-dependent feeding and identify candidate regulators of sugar-based food consumption in *Drosophila*.

## Introduction

Feeding is important for every organism’s survival, and it regulates biological processes including energy balance, lifespan (Bass et al., 2007; Skorupa et al., 2008), development (Layalle et al., 2005), and fecundity (Fricke et al., 2008). In humans, excessive consumption of high-sugar and processed foods has been linked to the rising incidence of type 2 diabetes, obesity, and cardiovascular disease (Baenas & Wagner, 2022; Goncalves et al., 2012). However, individuals exposed to the same foods do not develop these conditions at equal rates, which indicates that genetic background shapes how different individuals respond to the same food and that food and genotype interact to determine health outcomes (Francis et al., 2021). Identifying these variants underlying the interaction in humans is difficult, which motivates the use of *Drosophila melanogaster*, a genetically tractable model for our investigation.

The fruit fly is ideal for this study because fruit flies share conserved pathways for nutrient sensing, energy homeostasis, and metabolism with humans. Also, both food and genetic background affect fly metabolic phenotypes (Masek et al., 2014; Schwasinger-Schmidt et al., 2012). Feeding in fruit flies comprises a series of behaviors - foraging, sensory evaluation, meal initiation, ingestion, meal termination, and disengagement (Pool & Scott, 2014). Among these behaviors, consumption (the amount ingested during a feeding bout) is the primary determinant of nutrient intake and the trait we examined here.

Food consumption is not just controlled by nutrient availability. Flies regulate food intake to satisfy their caloric or nutritional demands rather than increasing intake as the nutrient level in food increases (Carvalho et al., 2005). Raising sugar or yeast concentration reduces total intake through compensatory feeding (Carvalho et al., 2005; Deshpande et al., 2014; Ja et al., 2009).

More generally, food intake is better explained by the balance of dietary protein and carbohydrate than by either alone (Bruce et al., 2013; Lee et al., 2008). Additionally, sustained exposure of flies to high-sugar blunts gustatory sensitivity and drives overconsumption (May et al., 2019), produces obesity-like phenotypes (Musselman et al., 2011; Skorupa et al., 2008), and induces thirst and dehydration (van Dam et al., 2020). Sugar-dependent consumption is thus a regulated, physiologically consequential trait whose regulation may vary genetically.

The *Drosophila* Genetic Reference Panel (DGRP) provides the appropriate resources to test if genetic diversity plays a role in regulating sugar-based food consumption. The DGRP comprises fully sequenced inbred lines derived from a natural population. It captures genetic variation and enables genome-wide association mapping of quantitative traits (Huang et al., 2014; Mackay et al., 2012). Previous studies have been conducted with DGRP to study genetic variation in food consumption. Garlapow et al. (2015) found significant genetic variation in mean food intake and its within-line variance, but this study was done on a single diet, 4% sucrose. So, the study resolves variation in consumption at one sugar level rather than variation in the response to different sugar levels. Francis et al. (2021) established gene-by-diet interaction in the DGRP across foods varying in sugar, fat, and protein, with the highest genetic variability on high-sugar foods. However, their phenotype was starvation resistance, not food consumption. Therefore, whether the compensatory food intake in response to different dietary sugar levels is under natural genetic control, and which loci govern it, remains unknown.

Variants identified by GWAS in the DGRP are predominantly non-coding and may be tags rather than causing the associated effect. Hits recovered at permissive significance thresholds can be false-positive, and the direction of an allelic effect in the DGRP does not always predict the effect of manipulating that gene in a common background. Therefore, functional validation is essential to confirm the effects of genes predicted by GWAS. Our GWAS identified 118 genes with variants at P < 10⁻⁵. We tested every gene for which viable knockdown flies could be obtained - 76 in total, the remainder having no available RNAi line or producing lethal knockdowns - using RNAi driven both ubiquitously and pan-neuronally, with intake measured under the same dietary conditions used for mapping.

In this study, we used GWAS of the DGRP to map genetic regulators of food consumption under low- and high-sugar foods. Flies adjust their intake in response to dietary sugar levels, and the extent of this adjustment varies among genotypes. The two dietary conditions recovered almost entirely separate sets of loci, indicating that consumption on each is largely under separate genetic control. RNAi knockdown of *CG42673* and *Transmembrane O-mannosyltransferase targeting cadherins 2 (Tmtc2)* reproducibly increased food intake, and knockdown of *M-spondin (mspo)* increased intake in most conditions tested. These genes are new starting points for understanding how sugar intake is controlled, and how energy balance is regulated more broadly.

## Materials and methods

### Drosophila stocks

All wild-type lines: *w^1118^*, Zimbabwe, Oregon-RC, Swedish-C, Samarkand, Urbana, Amherst, and Lausanne, 176 DGRP lines (Mackay et al., 2012), all GAL4 driver lines, all RNAi transgenic lines, and the corresponding control lines used in this study were obtained from Bloomington *Drosophila* Stock Center (BDSC). Flies were reared at 25°C under a 12 h:12 h light–dark cycle with 40-60% relative humidity on a standard cornmeal-based medium (Genesee Scientific, Cat#: 66-113) with the addition of 2% (w/v) yeast (Brewer’s yeast, MP Biomedicals). We used FlyBase to find information on phenotypes/function/stocks (Ozturk-Colak et al., 2024).

### Experimental diets

The experimental diets that were used for this study were made up of agar (Genesee Scientific, Cat#: 66-104), table sugar (sucrose), yeast (Brewer’s yeast, MP Biomedicals) and are referred to as “SY foods”. The sugar and yeast correspond to carbohydrate and protein respectively and different combinations of diets were made. We used 5% and 20% (w/v) sugar for the low and high respectively while the yeast level in the diets was fixed at 5% (w/v).

### Food consumption quantification

Food consumption was quantified with Con-Ex method (Shell & Grotewiel, 2022; Shell et al., 2021; Shell et al., 2018). For all food consumption assays, young, mated flies of the same sex and genotype (1-3 days old) were placed on experimental diets without dye for five days to acclimatize. At the end of the fifth day, five biological replicates consisting of 3-5 flies each were transferred to empty vials with the experimental foods mixed with 1% FD&C Blue #1 dye in a food cap. Flies were allowed to consume dyed food for 48 hours before terminating the experiment by transferring the flies into Eppendorf tubes for storage at -80°C.

The excreted dye from flies in the tubes was collected by vortexing in 3.5 mL of water, and its concentration was determined by absorbance at 630 nm. Optical density was extrapolated on a standard curve to calculate the volume of food consumed by flies. Statistical analyses were performed in GraphPad Prism (version 8). Because within-group variance scaled with the mean, consumption values were log-transformed before analysis. The data are presented on the original scale, and fold changes are ratios of geometric means. Comparisons between two groups used two-tailed Student’s t-tests, and comparisons across diet and genotype used two-way ANOVA, with Benjamini–Hochberg false discovery rate correction for multiple comparisons.

### Heritability of food consumption in DGRP

Broad-sense heritability (H²) of food consumption was estimated separately for each sex and each dietary condition using linear mixed models fitted by restricted maximum likelihood with the lme4 package in R [version 4.6.1]. For each sex-by-diet combination, food consumption was modelled as Food_consumption ∼ (1 | DGRP), with DGRP line as a random effect, and H² was calculated as σ²L / (σ²L + σ²E), where σ²L is the among-line variance component and σ²E the residual within-line variance. Ninety-five percent confidence intervals were obtained by parametric bootstrap with 1,000 resamples using bootMer. Heritability was not estimated for the low:high consumption ratio, which was calculated from line means and therefore has no within-line replication.

### Relationship between high and low sugar food consumption in DGRP

We computed Pearson’s product moment correlations for food consumption between high and low sugar diets in the DGRP lines.

### GWA analysis for food consumption

We performed GWA analyses using the 1,920,276 SNPs and indels with minor allele frequencies ≥ 0.05 on the DGRP webserver (Huang et al., 2014) (dgrp2.gnets.ncsu.edu).

Analyses were adjusted for the standard known covariates implemented by the server, namely Wolbachia infection status and the common polymorphic inversions, together with the line relatedness matrix. Covariate effects were tested by Type III ANOVA and are reported in Table S1. In(2R)NS was associated with male consumption on low-sugar food (*P* = 0.039) and Wolbachia infection status with the male consumption ratio (*P* = 0.018), while no covariate reached significance in the remaining four analyses.

We performed six analyses: male high sugar, male low sugar, ratio of low to high in male, female high sugar, female low sugar and ratio of low to high in female. The ratio trait was calculated for each line as the mean food consumption on the low-sugar food divided by mean consumption on the high-sugar food, computed separately within sex. This measure was used because absolute consumption differs substantially among lines, so a ratio captures the extent of compensatory adjustment independently of a line’s overall intake. Variants were considered candidates at *P* < 10⁻⁵ under the simple linear model or the mixed model, the threshold conventionally applied in DGRP association studies. Variants were assigned to genes using the annotation returned by the DGRP2 pipeline, yielding 118 candidate genes across the six analyses (Table 1). Gene ontology enrichment was performed in DAVID (Huang da et al., 2009a, 2009b) using FlyBase identifiers and the full annotated *Drosophila melanogaster* genome as background. Out of the 118 genes 96 genes were annotated for biological process, 101 for cellular component and 85 for molecular function. Terms were considered enriched at Benjamini–Hochberg adjusted *P* < 0.05, with nominally enriched terms reported separately (Table S2).

**Table 1.** Candidate genes associated with sugar-dependent food consumption in the DGRP. Variants reaching P < 1 × 10⁻⁵ under the simple linear model (Single P) or the mixed model (Mixed P) in at least one of six genome-wide association analyses, and the genes to which they map. MAF, minor allele frequency. The Analysis column indicates which association analyses recovered each gene: F, female; M, male; low, low-sugar food; high, high-sugar food; ratio, low:high consumption ratio. A total of 161 variants mapped to 118 genes.

| S/N | Gene name | FlyBase ID | Single P | Mixed P | MAF | SNP ID | Variant type | Analysis |
|---|---|---|---|---|---|---|---|---|
| 1 | *Als2* | FBgn0037116 | 2.16E-06 | 3.10E-06 | 0.06962 | 3L_21672453_SNP | NON_SYNONYMOUS_CODING | F high |
| 2 | *Ank2* | FBgn0261788 | 8.12E-06 | 2.07E-05 | 5.39E-02 | 3L_7701416_SNP | DOWNSTREAM, SYNONYMOUS_CODING | M ratio |
| 3 | *Argonaute-1* | FBgn0262739 | 1.12E-05 | 9.61E-06 | 0.4658 | 2R_9837340_SNP | INTRON | F high |
| 4 | *ATPase 8A* | FBgn0259221 | 1.59E-05 | 7.71E-06 | 1.69E-01 | 2R_9315880_SNP | INTRON | M ratio |
| 5 | *Attacin-A* | FBgn0012042 | 1.11E-05 | 9.30E-06 | 5.63E-02 | 2R_10636472_SNP | DOWNSTREAM, INTERGENIC | M ratio |
| 6 | *Attacin-B* | FBgn0041581 | 1.11E-05 | 9.30E-06 | 5.63E-02 | 2R_10636472_SNP | UPSTREAM | M ratio |
| 7 | *brat* | FBgn0010300 | 3.72E-07 | 3.46E-07 | 0.09615 | 2L_19139224_INS | INTRON | F low |
| 8 | *c(2)M* | FBgn0028525 | 1.18E-05 | 6.50E-06 | 0.3613 | 2L_16306491_SNP | DOWNSTREAM | M low |
| 9 | *Cad88C* | FBgn0038247 | 4.99E-06 | 2.71E-06 | 0.06173 | 3R_10465025_SNP | NON_SYNONYMOUS_CODING | F high |
| 10 | *cannonball* | FBgn0011569 | 3.96E-06 | 3.13E-06 | 0.1758 | 3L_10260948_SNP | UPSTREAM | M low |
| 11 | *Carbonic anhydrase 3* | FBgn0030056 | 1.08E-05 | 7.90E-06 | 1.63E-01 | X_8581141_SNP | INTRON | M ratio |
| 12 | *CG10254* | FBgn0027512 | 1.10E-05 | 8.31E-06 | 0.05455 | 3R_19519898_SNP | SYNONYMOUS_CODING | M high |
| 13 | *CG10561* | FBgn0002036 | 1.04E-05 | 8.45E-06 | 0.1538 | 2L_19116187_SNP | SYNONYMOUS_CODING | F low |
| 14 | *CG11030* | FBgn0031736 | 2.43E-06 | 1.38E-06 | 0.1026 | 2L_5755874_SNP | INTRON | F low |
| 15 | *CG11145* | FBgn0033168 | 9.56E-06 | 3.13E-05 | 0.4221 | 2R_3316120_SNP | UPSTREAM | M low |
| 16 | *CG11248* | FBgn0037117 | 2.16E-06 | 3.10E-06 | 0.06962 | 3L_21672453_SNP | UPSTREAM | F high |
| 17 | *CG1136* | FBgn0035490 | 9.11E-06 | 1.20E-05 | 0.239 | 3L_4057810_SNP | NON_SYNONYMOUS_CODING | F high |
| 18 | *CG12194* | FBgn0031636 | 3.07E-06 | 3.67E-06 | 5.13E-02 | 2L_4839164_DEL | UTR_5_PRIME | M ratio |
| 19 | *CG12499* | FBgn0038968 | 4.04E-06 | 3.23E-06 | 0.1049 | 3R_18271207_SNP | SYNONYMOUS_CODING | M high |
| 20 | *CG13579* | FBgn0035010 | 4.50E-06 | 5.48E-06 | 0.3007 | 2R_20365977_SNP | INTRON | F high |
| 21 | *CG13699* | FBgn0036781 | 2.04E-06 | 1.64E-06 | 4.37E-01 | 3L_18154012_SNP | INTRON | M ratio |
| 22 | *CG13901* | FBgn0035164 | 2.90E-06 | 1.52E-06 | 0.3145 | 3L_836430_SNP | DOWNSTREAM | M high |
| 23 | *CG13955* | FBgn0033412 | 1.91E-06 | 2.72E-06 | 6.88E-02 | 2R_5311411_SNP | INTERGENIC | M ratio |
| 24 | *CG14275* | FBgn0032022 | 8.68E-06 | 9.72E-06 | 0.3585 | 2L_8333457_SNP | SYNONYMOUS_CODING | M high |
| 25 | *CG14958* | FBgn0035413 | 5.28E-06 | 4.25E-06 | 0.2727 | 3L_3189651_SNP | UPSTREAM | F ratio |
| 26 | *CG14965* | FBgn0035414 | 5.28E-06 | 4.25E-06 | 0.2727 | 3L_3189651_SNP | DOWNSTREAM | F ratio |
| 27 | *CG15073* | FBgn0034379 | 2.48E-06 | 1.76E-06 | 0.05521 | 2R_14594891_SNP | SYNONYMOUS_CODING | F high |
| 28 | *CG15196* | FBgn0030296 | 7.05E-06 | 6.84E-06 | 0.4277 | X_11304656_SNP | NON_SYNONYMOUS_CODING | F high |
| 29 | *CG15594* | FBgn0037421 | 9.59E-06 | 1.02E-05 | 0.07453 | 3R_2116049_SNP | DOWNSTREAM, INTERGENIC | F high |
| 30 | *CG17349* | FBgn0032771 | 1.04E-05 | 9.31E-06 | 0.4762 | 2L_19372447_SNP | SYNONYMOUS_CODING | F ratio |
| 31 | *CG2444* | FBgn0030326 | 3.89E-06 | 4.23E-06 | 0.08537 | X_11597519_SNP | DOWNSTREAM | F high |
| 32 | *CG32373* | FBgn0052373 | 8.12E-06 | 2.07E-05 | 5.39E-02 | 3L_7701416_SNP | INTRON | M ratio |
| 33 | *CG32447* | FBgn0052447 | 8.79E-06 | 2.04E-05 | 0.1258 | 3L_21682978_SNP | DOWNSTREAM, INTERGENIC | M low |
| 34 | *CG33158* | FBgn0053158 | 6.30E-06 | 3.70E-06 | 0.1572 | 3L_16441921_SNP |  | M low |
| 35 | *CG34396* | FBgn0085425 | 3.51E-06 | 6.25E-06 | 4.11E-01 | 2R_16979026_SNP | INTERGENIC | M ratio |
| 36 | *CG42525* | FBgn0260430 | 5.28E-06 | 4.25E-06 | 0.2727 | 3L_3189651_SNP | NON_SYNONYMOUS_CODING | F ratio |
| 37 | *CG42553* | FBgn0260755 | 2.90E-06 | 1.52E-06 | 0.3145 | 3L_836430_SNP | DOWNSTREAM | M high |
| 38 | *CG42554* | FBgn0260756 | 2.90E-06 | 1.52E-06 | 0.3145 | 3L_836430_SNP | SYNONYMOUS_CODING | M high |
| 39 | *CG42673* | FBgn0261555 | 7.20E-07 | 1.57E-06 | 6.63E-02 | 3L_9560977_SNP | INTRON | M ratio |
| 40 | *CG42784* | FBgn0263354 | 2.77E-06 | 2.17E-06 | 0.3312 | 2L_13433914_SNP | INTRON | M low |
| 41 | *CG4301* | FBgn0030747 | 6.23E-06 | 6.28E-06 | 0.07273 | X_16305077_SNP | SYNONYMOUS_CODING | F high |
| 42 | *CG43083* | FBgn0262529 | 5.33E-06 | 1.17E-05 | 0.05988 | 3L_15643211_SNP | DOWNSTREAM | M low |
| 43 | *CG43084* | FBgn0262530 | 5.33E-06 | 1.17E-05 | 0.05988 | 3L_15643211_SNP | UPSTREAM | M low |
| 44 | *CG43980* | FBgn0264711 | 1.53E-06 | 2.36E-06 | 0.06918 | 3L_21651111_SNP | INTRON, UPSTREAM | F high |
| 45 | *CG44325* | FBgn0265413 | 5.88E-06 | 7.29E-06 | 1.29E-01 | X_8967942_INS | INTRON | M ratio |
| 46 | *CG5065* | FBgn0034145 | 5.50E-06 | 3.47E-06 | 1.46E-01 | 2R_12507214_SNP | INTRON | M ratio |
| 47 | *CG5861* | FBgn0015338 | 2.88E-06 | 1.16E-06 | 0.4324 | 2L_16307255_SNP | START_GAINED | M low |
| 48 | *CG5888* | FBgn0028523 | 9.31E-06 | 7.48E-06 | 0.1914 | 2L_16439822_SNP | INTRON | M high |
| 49 | *CG7886* | FBgn0038248 | 4.99E-06 | 2.71E-06 | 0.06173 | 3R_10465025_SNP | INTRON | F high |
| 50 | *CG9168* | FBgn0035216 | 2.95E-06 | 5.23E-06 | 9.52E-02 | 3L_1412526_SNP | SYNONYMOUS_CODING | F ratio; M ratio |
| 51 | *cni* | FBgn0000339 | 5.61E-06 | 2.68E-06 | 0.4675 | 2L_16308339_SNP | DOWNSTREAM | M low |
| 52 | *CR43903* | FBgn0264504 | 1.10E-05 | 8.31E-06 | 0.05455 | 3R_19519898_SNP | EXON | M high |
| 53 | *CR43979* | FBgn0264710 | 2.16E-06 | 3.10E-06 | 0.06962 | 3L_21672453_SNP | EXON | F high |
| 54 | *CR44203* | FBgn0265098 | 7.90E-06 | 2.52E-05 | 0.4323 | 2R_3316132_SNP | DOWNSTREAM | M low |
| 55 | *Ddc* | FBgn0000422 | 1.04E-05 | 8.45E-06 | 0.1538 | 2L_19116187_SNP | DOWNSTREAM | F low |
| 56 | *dlg1* | FBgn0001624 | 7.05E-06 | 6.84E-06 | 0.4277 | X_11304656_SNP | DOWNSTREAM | F high |
| 57 | *drop dead* | FBgn0260006 | 6.72E-06 | 6.75E-05 | 5.95E-02 | X_15024749_SNP | INTRON | M ratio |
| 58 | *dsx-c73A* | FBgn0261799 | 6.30E-06 | 3.70E-06 | 0.1572 | 3L_16441921_SNP | INTRON | M low |
| 59 | *Eag-like K+ channel* | FBgn0011589 | 8.62E-06 | 2.41E-05 | 0.07831 | 2R_13810884_SNP | INTRON | M low |
| 60 | *Ecdysone-induced protein 75B* | FBgn0000568 | 9.92E-06 | 5.68E-06 | 0.3289 | 3L_18021192_SNP | INTRON | F high |
| 61 | *egghead* | FBgn0001404 | 7.95E-06 | 8.14E-06 | 0.1104 | X_2492384_SNP | UTR_3_PRIME | F ratio |
| 62 | *Eig71Ea* | FBgn0004588 | 5.33E-06 | 1.17E-05 | 0.05988 | 3L_15643211_SNP | UPSTREAM | M low |
| 63 | *Eig71Eb* | FBgn0004589 | 5.33E-06 | 1.17E-05 | 0.05988 | 3L_15643211_SNP | NON_SYNONYMOUS_CODING | M low |
| 64 | *Eig71Ef* | FBgn0004593 | 3.09E-06 | 1.11E-05 | 0.06832 | 3L_15651307_SNP | INTERGENIC | M low |
| 65 | *Eig71Eg* | FBgn0004594 | 3.09E-06 | 1.11E-05 | 0.06832 | 3L_15651307_SNP | DOWNSTREAM | M low |
| 66 | *Eig71Eh* | FBgn0014848 | 3.09E-06 | 1.11E-05 | 0.06832 | 3L_15651307_SNP | UPSTREAM | M low |
| 67 | *enc* | FBgn0004875 | 7.08E-06 | 5.52E-06 | 2.63E-01 | 3L_3827300_SNP | INTRON | M ratio |
| 68 | *fizzy* | FBgn0001086 | 2.24E-06 | 2.58E-06 | 0.3525 | 2L_16311924_SNP | SYNONYMOUS_CODING | M low |
| 69 | *Glucose transporter 4 enhancer factor* | FBgn0263097 | 4.51E-06 | 2.85E-06 | 0.06369 | 3R_5792968_SNP | INTRON | M high |
| 70 | *Gpxl* | FBgn0034415 | 3.64E-05 | 9.03E-06 | 0.2901 | 2R_14943021_SNP | SYNONYMOUS_CODING | M low |
| 71 | *GUK-holder* | FBgn0026239 | 1.38E-06 | 9.54E-07 | 0.2628 | 3R_14814511_SNP | INTRON | F high |
| 72 | *homothorax* | FBgn0001235 | 4.01E-05 | 8.90E-06 | 0.1799 | 3R_6433122_INS | INTRON | M low |
| 73 | *Idit* | FBgn0053143 | 1.02E-05 | 7.37E-06 | 0.2548 | 2R_18511308_SNP | INTRON | F ratio |
| 74 | *Invadolysin* | FBgn0086359 | 1.23E-05 | 6.02E-06 | 0.2157 | 3R_5975217_SNP | INTRON | F low; M high |
| 75 | *julius seizure* | FBgn0039647 | 8.31E-06 | 3.80E-06 | 0.2635 | 3R_25008696_SNP | INTRON | F high |
| 76 | *Kank* | FBgn0027596 | 1.00E-05 | 9.66E-06 | 0.4214 | 2R_10804390_SNP | INTRON, SYNONYMOUS_CODING | F low |
| 77 | *kin of irre* | FBgn0028369 | 2.07E-06 | 8.84E-06 | 5.99E-02 | X_2713876_SNP | INTRON | M ratio |
| 78 | *Klp3A* | FBgn0011606 | 7.95E-06 | 8.14E-06 | 0.1104 | X_2492384_SNP | DOWNSTREAM | F ratio |
| 79 | *Liprin-α* | FBgn0046704 | 2.07E-06 | 7.28E-07 | 0.4684 | 2L_6729802_SNP | INTRON | M low |
| 80 | *long non-coding RNA:CR44231* | FBgn0265162 | 7.70E-06 | 7.96E-06 | 8.70E-02 | 3R_8554205_SNP | DOWNSTREAM, INTERGENIC | M ratio |
| 81 | *modifier of mdg4* | FBgn0002781 | 1.50E-05 | 7.52E-06 | 0.1329 | 3R_17196958_SNP | DOWNSTREAM, INTRON | F high |
| 82 | *Mothers against dpp* | FBgn0011648 | 1.53E-05 | 4.60E-06 | 0.3841 | 2L_3155669_DEL | INTRON | M low |
| 83 | *mRpL2* | FBgn0036135 | 4.95E-06 | 4.13E-06 | 0.135 | 3L_11115867_SNP | DOWNSTREAM | F ratio |
| 84 | *M-spondin* | FBgn0020269 | 2.49E-06 | 2.29E-06 | 0.07273 | 2R_10570521_SNP | INTRON | M low |
| 85 | *Myosuppressin receptor 1* | FBgn0035331 | 1.17E-05 | 8.60E-06 | 1.10E-01 | 3L_2325594_SNP | INTRON | M ratio |
| 86 | *Nadk1b* | FBgn0053156 | 6.16E-07 | 4.08E-07 | 0.2125 | 2R_9437990_SNP | INTRON | F low |
| 87 | *NaPi-III* | FBgn0260795 | 4.95E-06 | 4.13E-06 | 0.135 | 3L_11115867_SNP | UPSTREAM | F ratio |
| 88 | *ND-51L2* | FBgn0034007 | 1.09E-05 | 8.20E-06 | 0.06329 | 2R_11184249_SNP | DOWNSTREAM | F ratio |
| 89 | *nicotinic Acetylcholine Receptor α3* | FBgn0015519 | 1.18E-05 | 8.78E-06 | 0.09202 | X_8261368_SNP | UPSTREAM, INTERGENIC | F ratio |
| 90 | *patched* | FBgn0003892 | 5.68E-06 | 4.21E-06 | 2.83E-01 | 2R_4549044_SNP | SYNONYMOUS_CODING | M ratio |
| 91 | *Phosphorylase kinase γ* | FBgn0011754 | 1.18E-06 | 1.25E-06 | 0.07362 | X_11597539_SNP | INTRON | F high |
| 92 | *prel* | FBgn0033413 | 1.91E-06 | 2.72E-06 | 6.88E-02 | 2R_5311411_SNP | UPSTREAM | M ratio |
| 93 | *Protein tyrosine phosphatase 61F* | FBgn0003138 | 5.04E-06 | 4.07E-06 | 0.4323 | 3L_1433854_SNP | INTRON | F high; F ratio; M ratio |
| 94 | *PVR adaptor protein* | FBgn0052406 | 3.82E-06 | 4.04E-06 | 0.2112 | 3L_6008703_SNP | INTRON | F high |
| 95 | *Pykl5* | FBgn0037115 | 9.33E-06 | 1.62E-05 | 0.06369 | 3L_21664514_SNP | UPSTREAM, INTERGENIC | F high |
| 96 | *Rabaptin-5* | FBgn0034585 | 3.51E-06 | 6.25E-06 | 4.11E-01 | 2R_16979026_SNP | UPSTREAM | M ratio |
| 97 | *Red herring* | FBgn0064227 | 7.08E-06 | 5.52E-06 | 2.63E-01 | 3L_3827300_SNP | INTRON | M ratio |
| 98 | *Rgk3* | FBgn0085426 | 6.62E-06 | 5.76E-06 | 0.441 | 2R_17161035_SNP | INTRON | F low |
| 99 | *roadkill* | FBgn0264493 | 3.43E-06 | 6.62E-06 | 0.2115 | 3R_9833923_SNP | INTRON | F ratio |
| 100 | *SA-2* | FBgn0043865 | 5.04E-06 | 4.07E-06 | 0.4323 | 3L_1433854_SNP | DOWNSTREAM | F high |
| 101 | *Semaphorin 1a* | FBgn0011259 | 1.21E-05 | 8.20E-06 | 1.49E-01 | 2L_8660590_SNP | INTRON | M ratio |
| 102 | *Semaphorin 5c* | FBgn0250876 | 7.81E-06 | 1.12E-05 | 0.2516 | 3L_12067593_SNP | INTRON | F high |
| 103 | *SH2 ankyrin repeat kinase* | FBgn0015295 | 7.94E-06 | 8.24E-06 | 0.2405 | 2R_12089757_SNP | SYNONYMOUS_CODING | F ratio |
| 104 | *Shal K+ channel interacting protein* | FBgn0051163 | 8.15E-06 | 5.78E-06 | 0.07975 | 3R_18075434_SNP | INTRON | M high; M ratio |
| 105 | *sine oculis* | FBgn0003460 | 7.90E-06 | 2.52E-05 | 0.4323 | 2R_3316132_SNP | INTRON | M low |
| 106 | *small optic lobes* | FBgn0003464 | 9.98E-06 | 5.47E-06 | 0.4521 | X_21228258_DEL | INTRON | M high |
| 107 | *Sno oncogene* | FBgn0085450 | 4.99E-07 | 1.19E-06 | 1.37E-01 | 2L_7931651_SNP | INTRON | M ratio |
| 108 | *stumps* | FBgn0020299 | 6.13E-06 | 8.46E-06 | 0.2704 | 3R_10432113_SNP | SYNONYMOUS_CODING | M low |
| 109 | *Syndecan* | FBgn0010415 | 3.39E-06 | 7.85E-06 | 8.43E-02 | 2R_17293612_SNP | INTRON | M ratio |
| 110 | *Syntaxin 5* | FBgn0011708 | 5.61E-06 | 2.68E-06 | 0.4675 | 2L_16308339_SNP | INTRON | M low |
| 111 | *Tmtc2* | FBgn0028481 | 4.04E-06 | 5.84E-06 | 0.05521 | 2L_923173_SNP | UTR_5_PRIME | F high |
| 112 | *tonalli* | FBgn0026160 | 5.34E-06 | 9.69E-06 | 0.2795 | 3L_10857439_SNP | INTRON | M low |
| 113 | *Transient receptor potential cation channel, subfamily M* | FBgn0265194 | 1.09E-05 | 8.20E-06 | 0.06329 | 2R_11184249_SNP | INTRON | F ratio |
| 114 | *turtle* | FBgn0010473 | 8.15E-06 | 1.07E-05 | 5.99E-02 | 2L_4291011_SNP | INTRON | M ratio |
| 115 | *TwdlW* | FBgn0038487 | 8.70E-06 | 5.64E-06 | 0.2579 | 3R_12968721_SNP | UPSTREAM | F low |
| 116 | *Ubiquitin fusion-degradation 1-like* | FBgn0036136 | 4.95E-06 | 4.13E-06 | 0.135 | 3L_11115867_SNP | SYNONYMOUS_CODING | F ratio |
| 117 | *Ubiquitin specific protease 10* | FBgn0052479 | 8.94E-06 | 7.33E-06 | 0.473 | 3L_888574_SNP | SYNONYMOUS_CODING, UTR_5_PRIME | M high |
| 118 | *Vps51* | FBgn0034380 | 2.48E-06 | 1.76E-06 | 0.05521 | 2R_14594891_SNP | DOWNSTREAM | F high |

### Functional validation

Out of the 118 candidate genes identified, 76 were tested by RNAi knockdown, 25 had no available RNAi line, and knockdown of 17 was lethal. Three genes were represented by more than one independent line (*Shark*, three lines; *GUK-holder* and *PhKγ*, two each). Knockdown used the GAL4/UAS system with *arm*-GAL4 (ubiquitous) and *elav*-GAL4 (pan-neuronal). Sixty-eight genes were tested with both drivers, five were tested with *arm*-GAL4 only and four with *elav*-GAL4 only, because knockdown with the other driver produced no viable adults. Virgin females of each driver were crossed to males of each UAS-RNAi line at 25 °C. Because TRiP constructs are inserted at either attP2 or attP40, knockdown progeny were compared with progeny of the same driver crossed to the empty-vector control carrying the matched landing site (attP2, BDSC 36303; attP40, BDSC 36304).

Food consumption was measured by Con-Ex as described above, on the same low- and high-sugar foods used for mapping, with five biological replicates per genotype per diet. Within each combination of driver, sex and landing site, log-transformed consumption was analysed by two-way ANOVA with genotype and diet as factors, and each RNAi line compared with its matched control. *CG42673* and *Tmtc2* were retested in independent experiments using the same assay and controls, with *arm*-GAL4, *elav*-GAL4, *nSyb*-GAL4 and *repo*-GAL4.

## Results

### Dietary sugar levels regulate food consumption and compensatory feeding in *Drosophila*

Many previous studies have shown that food consumption in fruit flies changes in response to the level of nutrients in the food (Bruce et al., 2013; Carvalho et al., 2005; Chandegra et al., 2017; Shell et al., 2018). Flies eat less as carbohydrate and/or protein levels in the food increase (Beard et al., 2024; Deshpande et al., 2014; Ja et al., 2009). To explore how sugar (sucrose) affects food consumption, we measured consumption in the *w¹¹¹⁸* wild-type background on foods containing 5% (w/v) yeast and either 5% or 20% (w/v) sucrose, for low- and high-sugar foods, respectively. Adult male and female flies (1–3 days old) were maintained on the experimental foods without blue dye for five days, then transferred to the same foods containing 1% FD&C Blue #1 for 48 hours. Food intake was quantified by Con-Ex assay (Shell & Grotewiel, 2022; Shell et al., 2021; Shell et al., 2018), and volumes were extrapolated from a standard curve.

Flies consumed significantly more food on the low-sugar food than on high-sugar food, consistent across both sexes (males: 0.11 vs 0.06 μL/fly/day; females: 0.13 vs 0.07 μL/fly/day; two-tailed *t*-test, *P*<0.0001; Figure 1A). This indicates that flies adjust intake according to the caloric density of the food. This increase (on low sugar) and decrease (on high sugar) in food intake in response to the nutrient density of the food is referred to as compensatory feeding (Carvalho et al., 2005). Comparing sexes, females consumed more than males, particularly on the low-sugar food (two-way ANOVA, p=0.0402), while consumption did not differ significantly on the high-sugar food (two-way ANOVA, p=0.1852; Figure 1A). Because yeast content was identical across diets, greater intake of the low-sugar food also corresponds to greater protein intake, consistent with reported sexual dimorphism in macronutrient targets (Chandegra et al., 2017). Together, these results establish that dietary sugar concentration strongly influences food intake in flies and that overall consumption differs between sexes, particularly under low-sugar conditions. Flies consumed more food on low-sugar diets, reflecting compensatory feeding behavior in response to reduced sugar levels in the food.

**Fig. 1.**
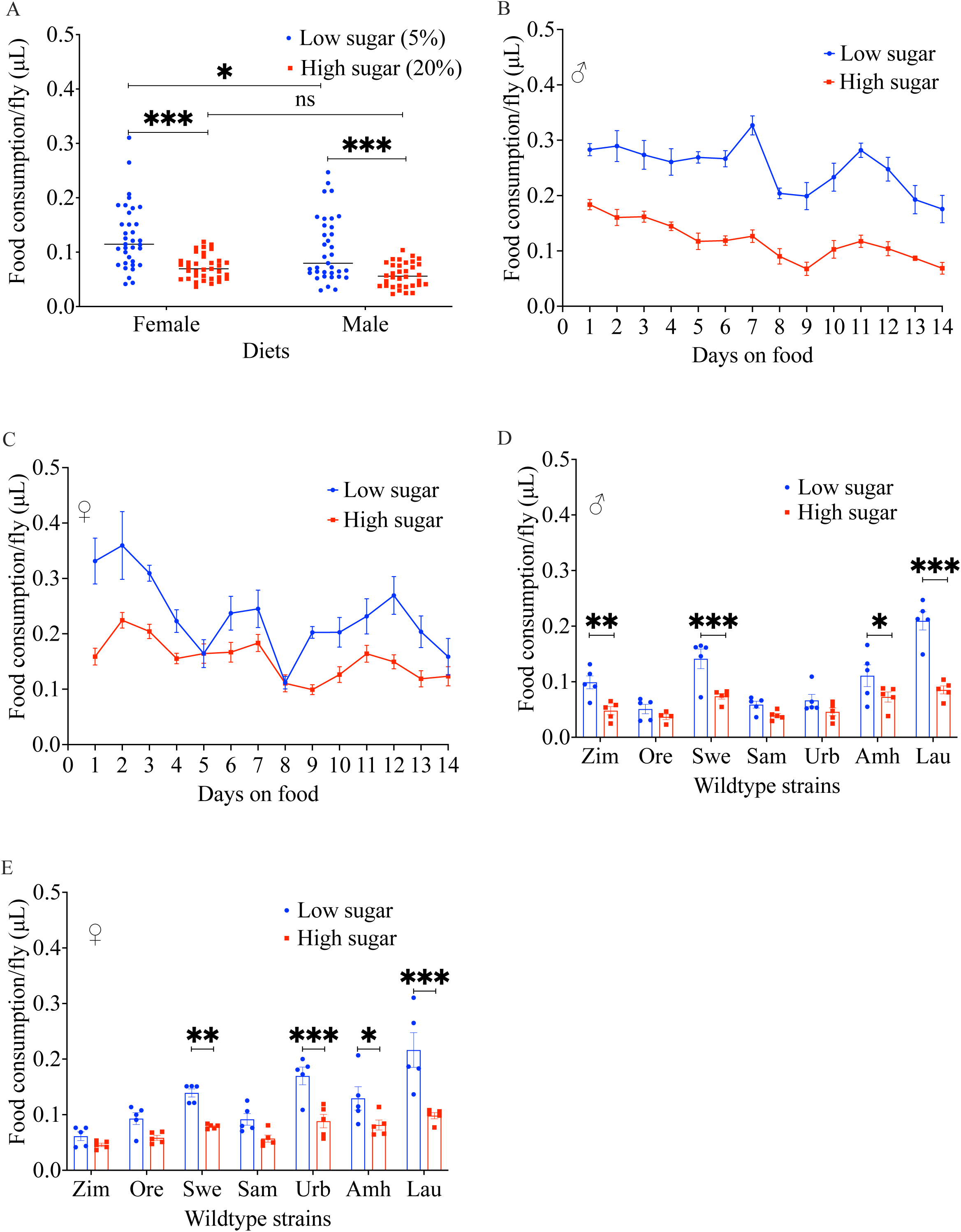
Dietary sugar levels influence food consumption and compensatory feeding behavior in *Drosophila melanogaster*. Flies were maintained on experimental foods containing 5% (w/v) yeast and either 5% (low) or 20% (w/v) sucrose (high). Flies were kept on the experimental foods without dye for five days, then transferred to the same foods containing 1% FD&C Blue #1. Food intake was quantified by Con-Ex assay and volumes extrapolated from a standard curve. Consumption is expressed as µL per fly per day. Flies were 1–3 days old at the start of each assay, and all experiments used five biological replicates of 3–5 flies per sex, genotype and diet. Data are mean ± SEM. (A) Food consumption of *w¹¹¹⁸* males and females on low- and high-sugar food. Both sexes consumed more on the low-sugar food. Females consumed more than males on the low-sugar food, with no significant difference on the high-sugar food. (B, C) Food consumption measured every 24 h for 14 days in *w¹¹¹⁸* males (B) and females (C) kept continuously on low- or high-sugar food. The same flies were followed throughout and transferred to fresh food daily. Males consumed more low-than high-sugar food at every time point. Females did so on all days except days 5 and 8, when consumption on the two foods did not differ. (D, E) Food consumption of seven wild-type strains in males (D) and females (E): Zimbabwe (Zim), Oregon-RC (Ore), Swedish-C (Swe), Samarkand (Sam), Urbana (Urb), Amherst (Amh) and Lausanne (Lau). Consumption differed significantly among strains, with Lausanne highest in both sexes and Oregon-RC (males) or Zimbabwe (females) lowest. Compensatory feeding was present in every strain, but its magnitude varied, from a low:high consumption ratio of 2.45 to 1.38 in males and 2.20 to 1.34 in females. Statistical comparisons used two-way ANOVA followed by Benjamini–Hochberg false discovery rate correction. Asterisks indicate adjusted P values: \**P* < 0.05, \*\**P* < 0.01, \*\*\**P* < 0.001, \*\*\*\**P* < 0.0001; ns, not significant.

### Compensatory feeding persists during prolonged exposure to high-sugar food

Prolonged exposure to high-sugar food changes sweet taste perception and promotes overconsumption in *Drosophila* (May et al., 2019). This finding raised the possibility that extended high-sugar feeding leads to loss of the compensatory feeding response observed on short-term exposure. To test whether long-term exposure to high-sugar food disrupts compensatory feeding, we measured food consumption every 24 hours for 14 days in flies maintained continuously on low- or high-sugar food. Flies were transferred to new food daily, and measurements were taken over 14 days on the same set of flies. In males, consumption on the low-sugar food surpassed that on the high-sugar food at every time point in the 14 days (Figure 1B; two-way ANOVA, effect of diet *P* < 0.0001). Females largely maintained compensatory feeding throughout, except on two days (Figure 1C; effect of diet *P* < 0.0001). Consumption on days 5 and 8 on low- and high-sugar food was not significantly different. These results imply that compensatory feeding is a behavior that persists in spite of continuous high-sugar exposure, rather than being gradually overturned as taste-adaptation alone would suggest. Having shown that the compensatory feeding response is consistent under prolonged sugar exposure, we next asked whether its magnitude varies with genetic background.

### Genetic background influences food intake and compensatory feeding

Multiple studies have reported compensatory feeding in different fly genotypes (Bruce et al., 2013; Carvalho et al., 2005; Chandegra et al., 2017; Shell et al., 2018). Whether the amount consumed and the magnitude of compensation vary with genetic background has not been examined. Therefore, we measured food consumption under the same low- and high-sugar conditions in seven wild-type laboratory strains derived from different geographical populations: Zimbabwe, Oregon-RC, Swedish-C, Samarkand, Urbana, Amherst, and Lausanne. Consumption varied considerably among strains. On the low-sugar food, Lausanne consumed the most in both sexes, four-fold more than Oregon-RC in males (least) and three-fold more than Zimbabwe in females (least) (Figure 1D, E). Lausanne also consumed the most on high-sugar food, twice the consumption of Oregon-RC in males (least) and of Zimbabwe in females (least). Two-way ANOVA revealed significant main effects of genotype (male: *F* = 23.07, df = 6, *P*<0.0001; female: *F* = 15.20, df = 6, *P*<0.0001) and sugar level (male: *F* = 67, df = 1, *P*<0.0001; female: *F* = 65.56, df = 1, *P*<0.0001), and a significant genotype × sugar interaction (male: *F* = 6.278, df = 6, *P*<0.0001; female: *F* = 3.541, df = 6, *P*=0.0048), indicating that the response to dietary sugar itself differs among genotypes.

Compensatory feeding was observed in every strain and in both sexes, but the magnitude varied several-fold. Compensatory coefficient - expressed as the ratio of low-sugar food consumed to high-sugar food consumed-ranged from 2.45 in Lausanne to 1.38 in Oregon-RC in males, and from 2.20 in Lausanne to 1.34 in Zimbabwe in females (Figure 1D, E). These findings show that genetic background shapes how much flies eat and how strongly they adjust intake to dietary sugar. Consistent with this, genetic variation in food intake has previously been reported among DGRP lines on a single sugar concentration (Garlapow et al., 2015). To identify the genetic variants underlying this variation, we next measured food consumption across lines of the *Drosophila* Genetic Reference Panel (DGRP).

### Food consumption varies widely across DGRP genotypes

Because the seven wild-type strains tested earlier showed substantial differences in food consumption across sugar foods (Figure 1D, E), we hypothesized that natural genetic variation contributes to differences in food intake. However, seven wild-type genotypes represent only a small fraction of the natural genetic diversity present in *Drosophila* populations. To survey variation more broadly, we measured food consumption across 176 lines from the *Drosophila* Genetic Reference Panel (DGRP) on both low- and high-sugar foods using Con-Ex method as described above (Shell & Grotewiel, 2022; Shell et al., 2021; Shell et al., 2018). Four lines yielded insufficient males and were excluded from male analyses (*n* = 172).

We observed substantial genetic variation in consumption (Figure 2A, B; p<0.0001). In males, DGRP-227 had the highest consumption on low sugar food, which was 16-fold more than DGRP-304, which had the least, while DGRP-517 had the highest consumption of high-sugar food, which was 15-fold more than DGRP-306, which had the least. In females, DGRP-808 had the highest consumption of low-sugar food, which was almost 8-fold more than DGRP-217, which had the least, while DGRP-517 had the highest consumption of high-sugar food, which was 8-fold more than DGRP-783, which had the least. Two-way ANOVA revealed significant effects of genotype (male: *F* = 14.54, df = 171, *P*<0.0001; female: *F* = 9.714, df = 175, *P*<0.0001), and sugar level (male: *F* = 2018, df = 1, *P*<0.0001; female: *F* = 1662, df = 1, *P*<0.0001), a significant genotype × sugar interaction (male: *F* = 6.027, df = 171, *P*<0.0001; female: *F* = 3.462, df = 175, *P*<0.0001), indicating that the response to dietary sugar differs among the DGRP genotypes as it did among wildtype strains. Genetic variation in food intake among DGRP lines has been reported previously on a single sugar concentration (Garlapow et al., 2015). Our study extends this to two sugar concentrations (low and high) and shows that genotypes differ in how they respond to the difference in sugar levels. Because compensatory feeding was observed across the wild-type strains, we next asked whether this behavior is also maintained across the broader genetic diversity of the DGRP.

**Fig. 2.**
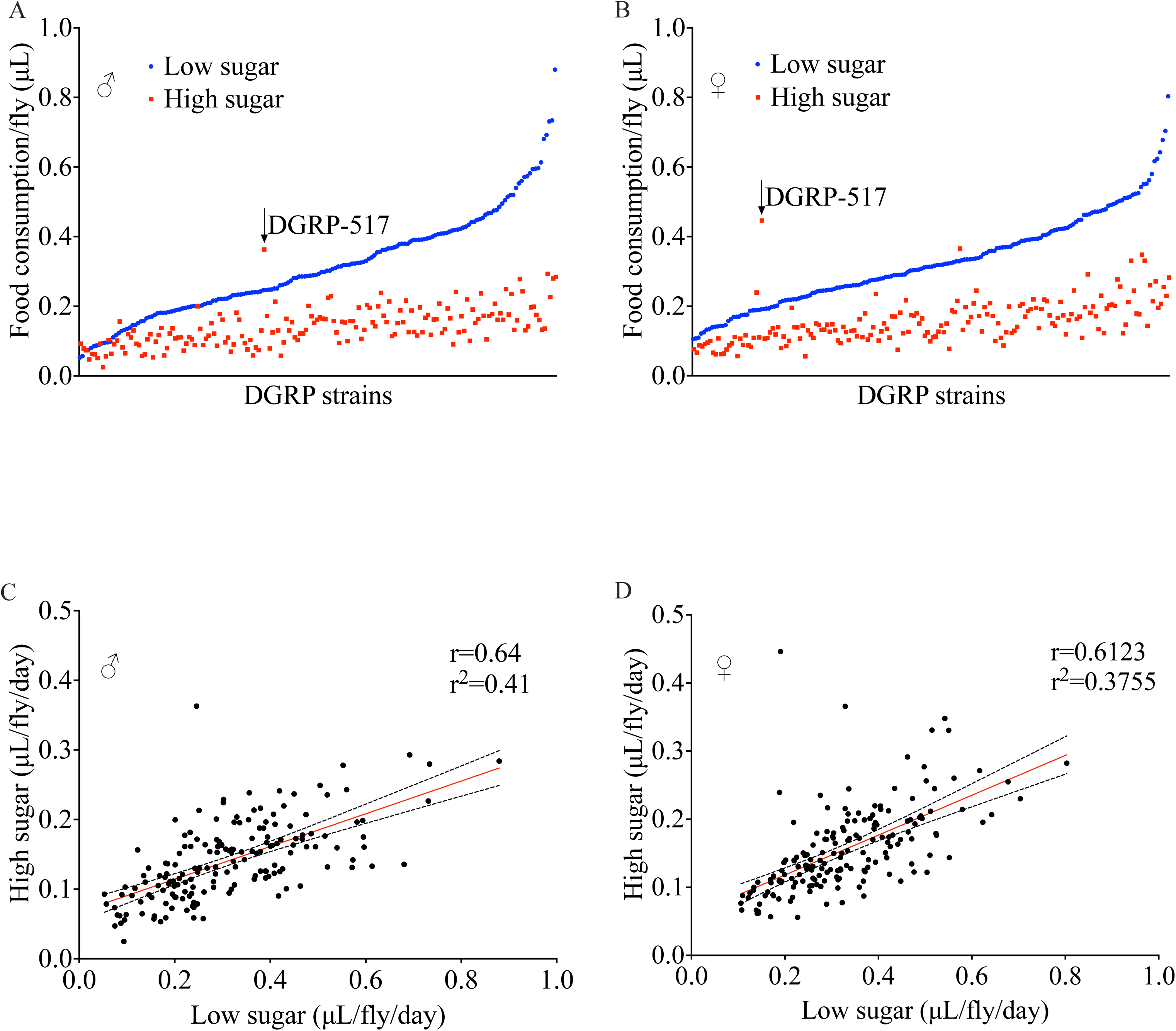
Extensive genetic variation in food consumption and compensatory feeding across the *Drosophila* Genetic Reference Panel (DGRP). Food consumption was measured in 176 DGRP lines (172 for males) on foods containing 5% (w/v) yeast and either 5% (low) or 20% (w/v) sucrose (high). Flies were 1–3 days old at the start of the assay and were kept on the experimental foods without dye for five days, then transferred to the same foods containing 1% FD&C Blue #1 for 48 h. Intake was quantified by the Con-Ex assay and volumes extrapolated from a standard curve. Consumption is expressed as µL per fly per day, from five biological replicates of 3–5 flies per line, sex and diet. The same line means shown in stock order appear in Figure S1. (A, B) Line mean consumption in males (A) and females (B), with lines ordered by consumption on the low-sugar food. Consumption on the low-sugar food exceeded that on the high-sugar food in 173 of 176 lines. Of the three exceptions, only DGRP-517 (arrow) reached significance after correction for multiple testing. (C, D) Correlation between line mean consumption on low- and high-sugar food in males (C) and females (D). Consumption on the two foods was positively correlated (males: r = 0.64, r² = 0.41; females: r = 0.61, r² = 0.37; both *P* < 0.0001). Solid lines show the linear regression and dotted lines its 95% confidence band. Comparisons in a and b used two-way ANOVA followed by Benjamini–Hochberg false discovery rate correction. Correlations in C and D are Pearson’s product-moment correlations computed on line means.

### *Drosophila* maintains compensatory feeding in the DGRP population

Studies have shown that fruit flies compensate for increased nutrient by reducing the quantity of food consumed (Beard et al., 2024; Carvalho et al., 2005; Deshpande et al., 2014), and our result (Figure 1D, E) showed that different sugar levels in food also make fruit flies exhibit this phenotype. We also observed that the degree of compensation varies among genotypes, which suggests that different genotypes differ in how strongly they adjust intake. We therefore asked whether any naturally derived genotypes instead consume more high-than low-sugar food.

To test this hypothesis, we compared food consumption of DGRP flies on low- and high-sugar foods with Benjamini–Hochberg correction applied across the full panel. Interestingly, consumption on the low-sugar food exceeded that on the high-sugar food in 173 of 176 lines (Figure 2A, B). One line, DGRP-517, consumed significantly more of the high-sugar food, and the other two showed a non-significant trend in the same direction. The near-complete absence of reversals across the 176 DGRP lines indicates that compensatory feeding response is a highly conserved feature of feeding regulation in this population, whether because the trait is under selection or because intake at high nutrient density is physiologically constrained. While compensatory feeding remained largely conserved, the amount of food consumed varies widely among lines on both foods. To further understand the relationship between feeding across different sugar levels, we examined whether food intake on low- and high-sugar foods is correlated across genotypes.

### Food consumption on low- and high-sugar foods is positively correlated

We found a strong positive correlation between the amount of food consumed on low- and high-sugar foods (males: r = 0.6406, p<0.0001; females: r = 0.6128, p<0.0001; Figure 2C and 2D), indicating that lines with higher baseline consumption tend to consume more regardless of sugar concentration. The correlation only accounted for 41% of the variance in males and 37% in females, so most of the variation in consumption is not explained by the consumption on the other food. Much of the variation therefore depends on which food a line is eating, not simply on how much that line eats overall. The extensive variation in food consumption observed across DGRP lines suggests that genetic factors contribute significantly to this trait. To quantify the contribution of genetic variation, we estimated the broad-sense heritability of food consumption on low- and high-sugar foods.

### Sugar-dependent food consumption is a heritable trait in the DGRP population

Broad-sense heritability helps determine the extent to which phenotypic variation is explained by genetic differences and whether genome-wide association studies (GWAS) are likely to identify associated loci (Visscher et al., 2008; Zaitlen & Kraft, 2012). Given the variation we observed in low- and high-sugar food consumption, we estimated heritability (H²) for each food and sex using linear mixed models with DGRP line as a random effect. Heritability was substantial in both sexes and on both foods. In males, H² was 0.66 (95% CI 0.60–0.71) on the low-sugar food and 0.62 (0.55–0.68) on the high-sugar food. In females, the corresponding estimates were 0.53 (0.46–0.59) and 0.56 (0.49–0.62). Estimates were somewhat higher in males than females, though the confidence intervals overlap. Genetic differences among lines therefore account for more than half of the variation in food consumption on both foods, indicating that sugar-dependent food consumption is strongly influenced by genetic background and well suited to genome-wide association mapping. Having established that sugar-based food consumption is substantially heritable, we used genome-wide association analysis to identify the variants contributing to this variation.

### GWAS identifies genetic variants associated with sugar-based food consumption

Given the substantial heritability of sugar-based food consumption, we used GWAS to identify genetic variants associated with variation in food intake under low- and high-sugar conditions. A previous study identified novel genes affecting food intake using GWAS (Garlapow et al., 2015) with a single sugar concentration. Here, we analyzed each dietary condition separately as well as the ratio between them. We performed six independent analyses using the DGRP2 pipeline: consumption on low-sugar food, consumption on high-sugar food, and a food consumption coefficient, defined as the ratio of intake on low- to high-sugar food, each in males and females separately. Phenotypes were adjusted for Wolbachia infection status and the five common polymorphic inversions. Covariate effects were generally small (Table S1): In(2R) NS was associated with male consumption on the low-sugar food (*P* = 0.039) and Wolbachia infection status with the male consumption ratio (*P* = 0.018), while no covariate reached significance in the remaining four analyses. All analyses were adjusted for the full covariate set regardless of significance.

Candidate variants identified across the six GWAS analyses were combined to generate a comprehensive list of loci associated with sugar-based food consumption. At a nominal significance threshold of *P* < 1 × 10⁻⁵ under the simple linear or mixed model, we found 161 SNPs significantly associated with sugar-based food consumption. These SNPs were mapped to 118 candidate genes, based on the combined results from all six GWAS analyses (Figure 3A-F, Table 1). The number of associated variants differed considerably between analyses: 21 and 11 for male consumption on low- and high-sugar food, 25 and 42 for the corresponding female analyses, and 49 and 13 for the male and female consumption ratios. The male ratio analysis yielded both the largest number of associated variants and the strongest single association in the study (*P* = 6.2 × 10⁻⁸), and the two ratio analyses together contributed 62 of the 161 variants. The compensatory response is therefore associated with genetic variation at least as extensively as consumption itself. Notably, no variant was shared between analyses, and only four genes were recovered by more than one: *Ptp61F*, associated with female high-sugar consumption and with the consumption ratio in both sexes; *CG9168* and *SKIP*, each recovered twice; and *Invadolysin*, associated with female low-sugar and male high-sugar consumption. Consumption on the two foods and the ratio between them therefore map to almost entirely distinct sets of loci, consistent with the limited correlation observed between intake on the two diets. Because only 176 lines were assayed, genome-wide significance (*P* = 2.6 × 10⁻⁸ after Bonferroni correction for 1.9 million variants) is unattainable for variants of realistic effect size, and a threshold of *P* < 1 × 10⁻⁵ is conventional in DGRP studies. This threshold admits false positives, so associated genes were treated as candidates to be tested experimentally rather than as established associations. Therefore, all 118 were carried forward to functional validation.

**Fig. 3.**
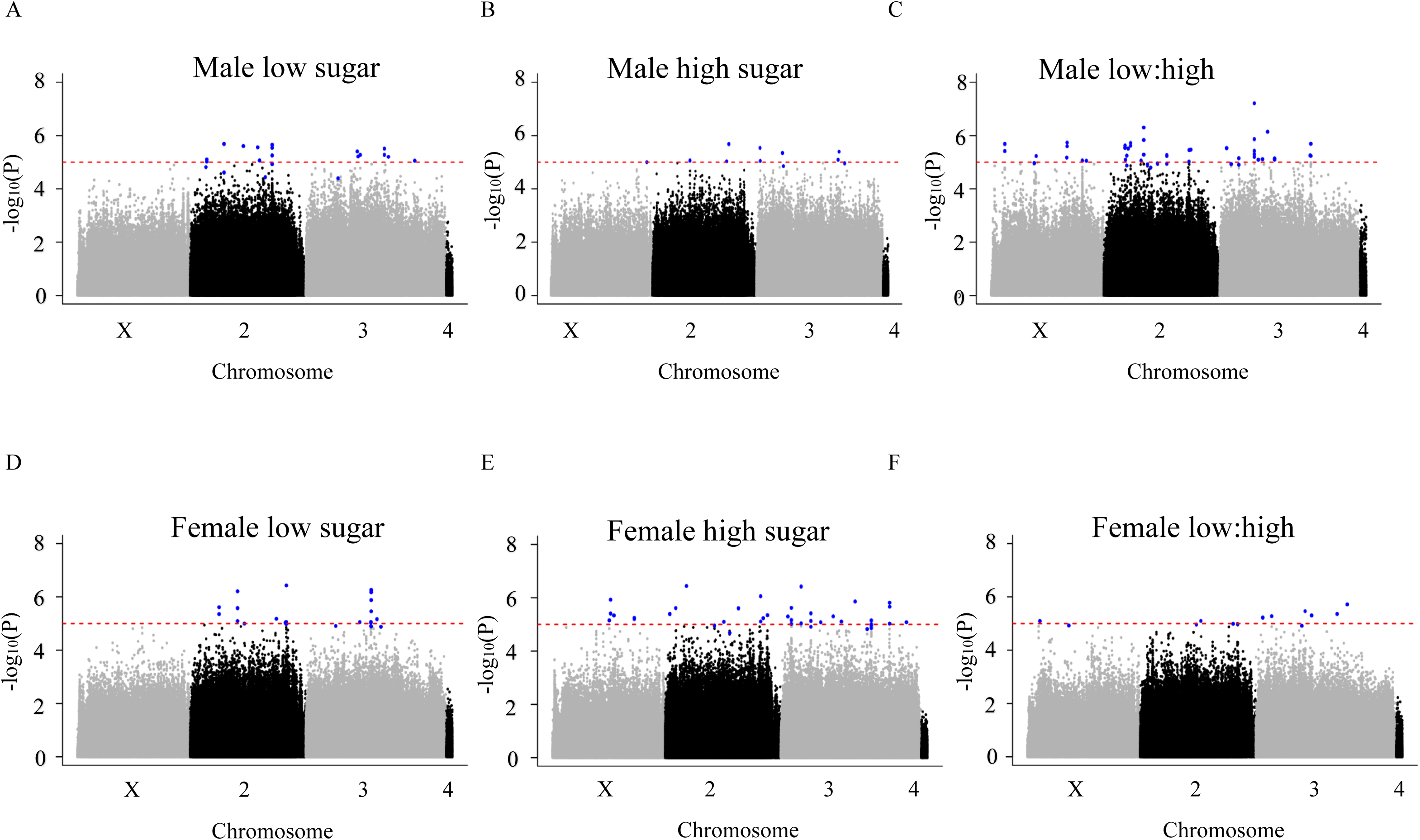
Genome-wide association analysis identifies candidate loci associated with sugar-based food consumption in *Drosophila*. Associations were tested for 1,920,276 SNPs and indels with minor allele frequency ≥ 0.05 using the DGRP2 pipeline, in 176 DGRP lines (172 for males) maintained on foods containing 5% (w/v) yeast and either 5% or 20% (w/v) sucrose. Six analyses were performed: consumption on low-sugar food, consumption on high-sugar food, and the low:high consumption ratio, each in males and females. Manhattan plots show −log₁₀(P) against genomic position across chromosomes X, 2, 3 and 4, with alternating grey and black indicating successive chromosomes. (A-C) Male analyses: consumption on low-sugar food (A), on high-sugar food (B), and the low:high consumption ratio (C). (D-F) Female analyses: consumption on low-sugar food (D), on high-sugar food (E), and the low:high consumption ratio (F). Blue points mark variants reaching *P* < 1 × 10⁻⁵ under the simple linear model or the mixed model. The dashed red line marks *P* = 1 × 10⁻⁵ for the simple linear model, which is the value plotted on the y-axis; blue points falling below this line were significant under the mixed model only. Across the six analyses, 161 variants exceeded the threshold and mapped to 118 candidate genes (Table 1).

Gene ontology enrichment analysis of the 118 candidate genes was performed in DAVID against the full annotated *Drosophila melanogaster* genome. One term, defense response to bacterium, was enriched after Benjamini–Hochberg correction (7 genes, 11.5-fold, *P* = 2.9 × 10⁻⁵, Benjamini = 1.3 × 10⁻², Table S2). However, this signal derives from two tandem gene clusters — *Attacin*-*A* and *Attacin-B*, tagged by a single shared variant, and five adjacent Eig71E genes — rather than from independently associated loci, and therefore reflects the physical clustering of antimicrobial peptide genes rather than genuine functional enrichment. Remaining significant terms described sequence-composition features (disordered and low-complexity regions) that track protein length rather than function. At nominal *P* < 0.05, the most enriched biological categories related to synaptic organization and neuronal function, including presynaptic membrane, synaptic assembly at the neuromuscular junction, axon guidance, and regulation of membrane potential. None survived correction for multiple testing. Therefore, no enrichment survived correction with a clear biological interpretation. This fits a polygenic trait, where many genes across unrelated processes each contribute a small amount, rather than one pathway driving the variation. These results identify a set of candidate loci potentially involved in regulating sugar-based food consumption in *Drosophila.* To determine whether these candidate genes directly influence sugar-based food consumption, we performed functional validation with RNAi knockdown.

### RNAi knockdown reveals candidate regulators of sugar-based feeding

Not all genes identified by GWAS are expected to directly regulate sugar-based food consumption because associations at *P* < 10⁻⁵ include false positives. Therefore, we performed functional validation using RNAi-mediated knockdown via the UAS-GAL4 system with TRiP lines from BDSC. Of the 118 candidate genes, 76 could be tested, knockdown of 17 was lethal during developmental stage, and 25 had no available RNAi line. TRiP constructs are inserted at either attP2 or attP40, so we compared each line with the empty-vector control at its landing site. To examine the global effect of each gene, we first performed ubiquitous knockdown. The initial trial using the strong ubiquitous driver, actin-GAL4, resulted in developmental lethality in most of the RNAi lines. To minimize lethality associated with strong ubiquitous knockdown, we switched to the weaker ubiquitous driver, *arm*-GAL4. Across all knockdown lines, flies consumed more low-sugar food than high-sugar food in both sexes (Figure S2), consistent with compensatory feeding observed in wild-type flies. Consumption relative to controls, however, varied widely (Figure 4A, B). In females, 29 of 77 lines altered consumption on the low-sugar food (22 increased, 7 decreased) and 19 on the high-sugar food (12 increased, 7 decreased). In males, 22 of 74 lines altered consumption on the low-sugar food (10 increased, 12 decreased) and 13 on high-sugar food (5 increased, 8 decreased).

**Fig. 4.**
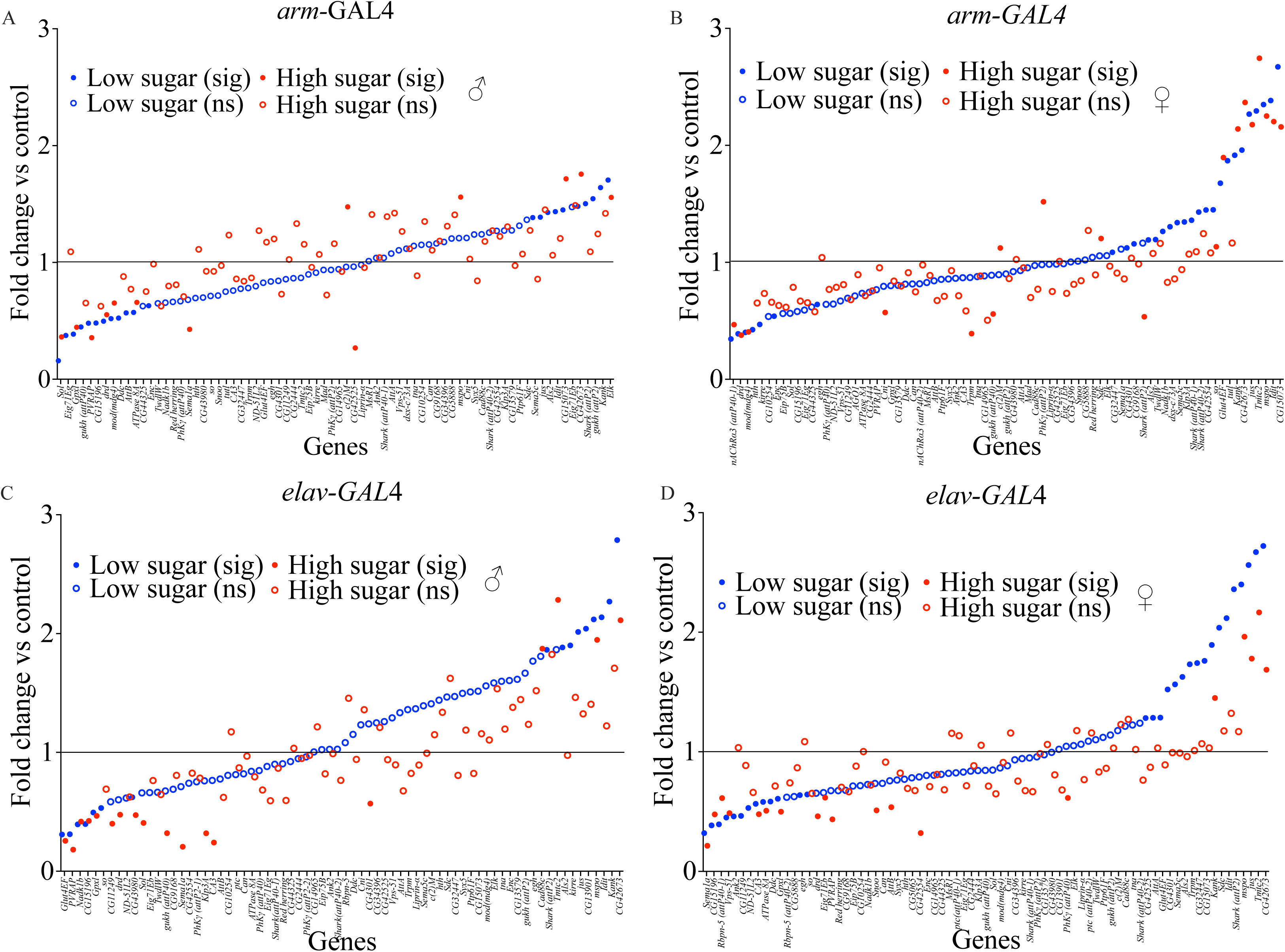
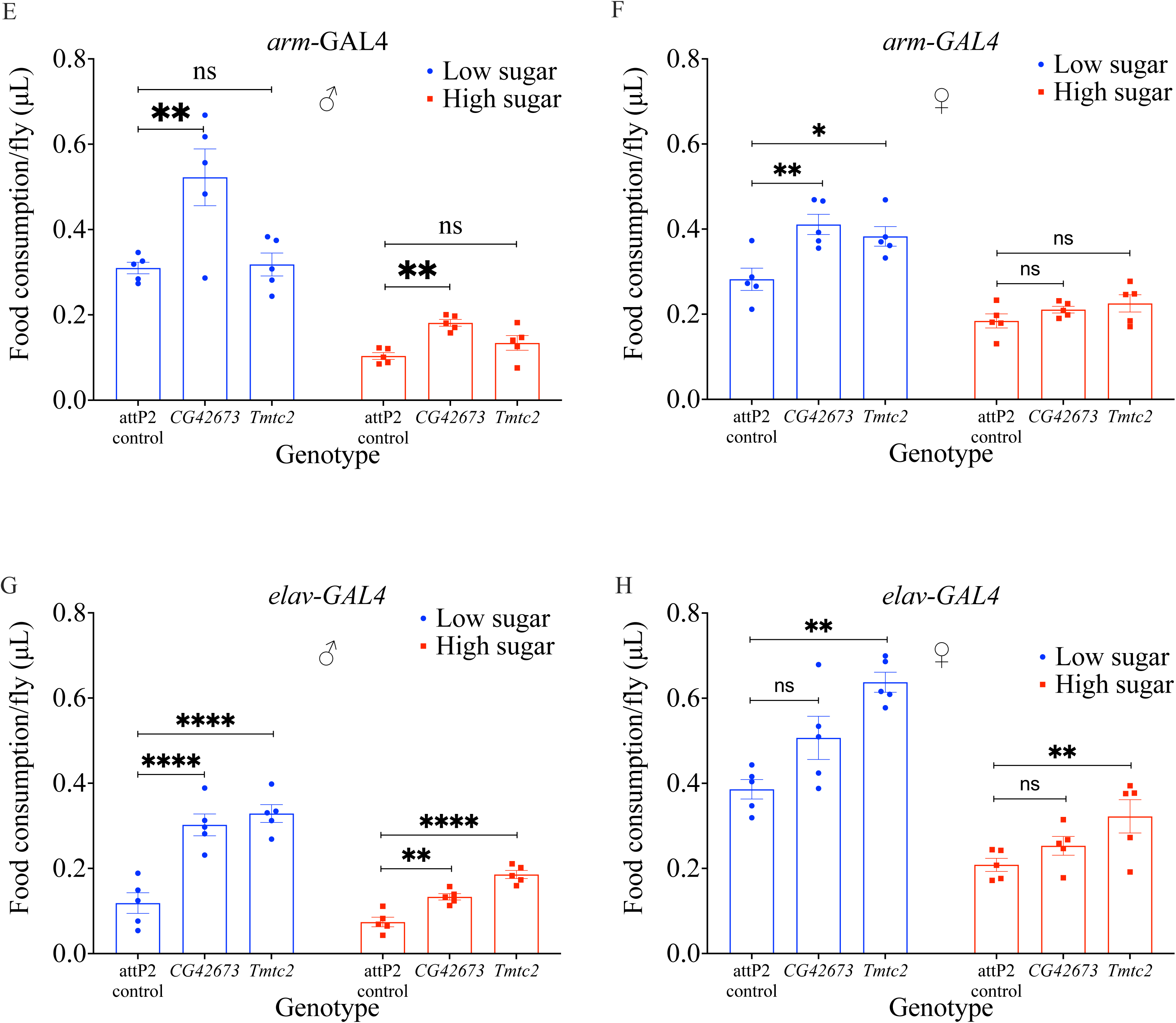
RNAi functional validation identifies candidate regulators of sugar-based food consumption in *Drosophila melanogaster*. Functional validation of candidate genes identified from GWAS was performed using RNAi-mediated knockdown via the UAS–GAL4 system. RNAi constructs were inserted at one of two genomic landing sites (attP2 or attP40). RNAi lines were grouped according to their insertion site and compared with the corresponding site-matched control line (attP2, BDSC 36303; attP40, BDSC 36304). Food consumption was measured using the Con-Ex assay on foods containing 5% yeast (w/v) and either low sugar (5% sucrose, w/v) or high sugar (20% sucrose, w/v). Data represent 5 biological replicates per genotype with 3–5 flies per replicate. (A–D) Effects of RNAi knockdown on food consumption relative to insertion-site controls, following ubiquitous knockdown with *arm*-GAL4 in males (A) and females (B), and pan-neuronal knockdown with *elav*-GAL4 in males (C) and females (D). Each point is one RNAi line, ordered within each insertion-site block by fold change on low-sugar food. Filled symbols indicate a significant change relative to control, open symbols no significant change. The long horizontal black line indicates the control baseline. Across both drivers, knockdown of 57 of 76 genes altered consumption in at least one condition. (E–H) Independent confirmatory experiment for *CG42673* and *Tmtc2*, showing food consumption following ubiquitous knockdown with *arm*-GAL4 in males (E) and females (F), and pan-neuronal knockdown with *elav*-GAL4 in males (G) and females (H). Knockdown of both genes reproducibly increased food intake, with the largest effects under pan-neuronal knockdown in males. Consumption values were log-transformed before analysis to equalize variance between the two sugar concentrations. For (A–D), significance was determined using two-way ANOVA followed by Benjamini–Hochberg false discovery rate (FDR) correction applied within each driver, sex and insertion-site. For (E–H), significance was determined using two-way ANOVA followed by Dunnett’s multiple comparisons test against the attP2 control within each sugar concentration. Asterisks indicate adjusted P values: \**P* < 0.05, \*\**P* < 0.01, \*\*\**P* < 0.001, \*\*\*\**P* < 0.0001, ns, not significant.

While *arm*-GAL4 drives ubiquitous expression, tissue-specific GAL4 drivers allow us to identify the tissues responsible for feeding regulation. Given the nervous system’s central role in appetite and satiety control, we performed pan-neuronal knockdown using *elav*-GAL4 to determine whether these genes’ effects were mediated through the nervous system (Figure 4C, D; Figure S3). In females, 30 of 77 lines altered consumption on the low-sugar food (17 increased, 13 decreased) and 19 on the high-sugar food (5 increased, 14 decreased). In males, 16 of 72 lines altered consumption on the low-sugar food (9 increased, 7 decreased) and 18 on the high-sugar food (4 increased, 14 decreased). Overall, 57 of the 76 genes across both drivers affected consumption in at least one condition, with increases and decreases about equally common. A small number of genes acted consistently across the eight combinations of driver, sex and diet. Knockdown of *CG42673* increased consumption in all eight (1.6- to 3.0-fold), *mspo* in seven, *Kank* and *jus* in six, and *Tmtc2*, *CG15073* and *Idit* in five. *Tmtc2* knockdown increased consumption strongly under both drivers in females and under *elav*-GAL4 in males on high-sugar food but had no effect under *arm*-GAL4 in males. Conversely, knockdown of *drop dead* and *Gpx-like* consistently decreased consumption (six conditions each).

To confirm the strongest effects, *CG42673* and *Tmtc2* were retested with both drivers in an independent experiment (Figure 4E–H). Ubiquitous knockdown again increased food consumption. *CG42673* knockdown increased food intake in males on both sugar foods (1.6-fold on low sugar, *P* = 0.004, and 1.8-fold on high sugar, *P* = 0.001) and in females on low sugar (1.5-fold, *P* = 0.003). *Tmtc2* knockdown increased intake in females on low sugar (1.4-fold, *P* = 0.012) but had no effect in males on either sugar food (1.0- and 1.3-fold, P ≥ 0.20), reproducing the sex difference seen in the primary screen. Pan-neuronal knockdown had larger effects but was less consistent. Under *elav*-GAL4 both genes increased male consumption on both sugar foods (1.9- to 3.0-fold, P ≤ 0.003), and *Tmtc2* increased female consumption on both sugar foods (1.5- and 1.7-fold, P ≤ 0.007). In males these increases were similar in the primary screen and the confirmatory experiment (2.4- and 2.3-fold for *CG42673*, 1.9- and 2.8-fold for *Tmtc2*). In females they were not consistent, ranging from a 2.6-fold increase to a 0.7-fold reduction across four independent experiments (Figure 4D, H and Supplementary Figure 4A, B). A second pan-neuronal driver, *nSyb*-GAL4, gave no consistent effect across two independent trials (Supplementary Figure S4C, D). Of eight comparisons only one reached significance (*Tmtc2* on low sugar, 1.7-fold, *P* = 0.012) and it did not replicate in the second trial (1.1-fold, *P* = 0.88).

Glial knockdown with *repo*-GAL4 instead reduced consumption in every comparison (0.4-fold on low sugar, 0.5- to 0.6-fold on high sugar, P ≤ 0.028, Supplementary Figure S4E), a reduction large enough that it may reflect impaired general health rather than a specific effect on feeding. Together, these results show that genes identified by mapping natural variation in food consumption alter consumption when knocked down directly. Ubiquitous knockdown of *CG42673* increased intake in males in two independent experiments and on both sugar foods, and increased intake in females on the low-sugar diet. Knockdown of *Tmtc2* increased intake in females, also on the low-sugar diet. These effects suggest that both genes normally act to limit consumption, although the tissue in which they act remains unresolved.

## Discussion

### Compensatory feeding on sugar diet

Flies of both sexes reduced intake as dietary sugar increased, consistent with compensatory feeding in response to nutrient density (Carvalho et al., 2005; Ja et al., 2009; Shell et al., 2018). The most striking feature of this response across our experiments was the stability of its direction. Compensation persisted over 14 days of continuous exposure, in all seven wild-type strains, in 175 of 176 DGRP lines, and in every line of the ubiquitous RNAi screen.

However, its magnitude varied several-fold among genotypes regardless of sex, from 34% to 145% greater intake on the low-sugar food across the wild-type strains. Whether flies compensate appears to be fixed. How strongly they compensate is not, and it was this variation that we set out to map.

Because flies in our assay had access to only one food at a time, the compensation we measured reflects regulation of how much is eaten rather than a choice between foods. Flies do have mechanisms that pull them toward caloric sugars, including a calorie-induced secreted factor released after ingestion (Abu et al., 2018) and learned associations with post-ingestive reward (Burke & Waddell, 2011). Both should have increased consumption of the high-sugar food, yet flies ate less of it. This suggests that when there is no choice to make, internal caloric signals appear to outweigh palatability. The same reasoning applies to the sexes. Males show a stronger preference for carbohydrate than females in choice assays (Chandegra et al., 2017; Diegelmann et al., 2017), which would predict weaker compensation in males, yet males maintained compensation throughout the 14-day exposure while intake in females converged on days 5 and 8. A more likely explanation for the female pattern is their changing nutritional demand during egg production (Baker et al., 2007; Barnes et al., 2008). Because yeast content was identical in both diets, eating more of the low-sugar food also delivers more protein, so the protein requirements of mated females may override caloric regulation from time to time.

Although most studies report compensatory feeding in *Drosophila*, some report the opposite, with flies eating more as sugar concentration increases (Diegelmann et al., 2017; Edgecomb et al., 1994; May et al., 2019; Min & Tatar, 2006). These differences may reflect assay design as much as biology. Diegelmann et al. offered flies a simultaneous choice among five sucrose concentrations over three hours on sugar-only liquid food (Diegelmann et al., 2017). Given a choice, flies take the most caloric option. Compensation shows up under different conditions: one food, and enough time for post-ingestive feedback to act. Differences in quantification method (Min & Tatar, 2006), diet composition and assay duration are therefore likely contributors, as is genetic background. The several-fold variation in compensation we observed among genotypes suggests that laboratories using different strains may obtain genuinely different results from the same protocol.

### Variation in food consumption

Food intake varied substantially among genotypes, in both the wild-type strains and the DGRP. Lausanne females consumed roughly 3.5 times more low-sugar food than Zimbabwe females, and among DGRP lines male intake on low-sugar food spanned about 16-fold. This is consistent with earlier reports of extensive natural variation in metabolic and behavioural traits in the panel (Francis et al., 2021; Garlapow et al., 2015; Jehrke et al., 2018; Unckless et al., 2015). Much of this noticed variation is due to genetics. Broad-sense heritability of consumption ranged from 0.53 to 0.66 across sexes and diets, higher than the 0.45 reported for food intake on a single sucrose food (Garlapow et al., 2015). Genetic differences between lines therefore account for more than half the variation in how much a fly eats, which is enough for association mapping.

However, high heritability does not mean the trait is controlled by a few genes. The associated variants were scattered across the genome rather than concentrated in a few regions, so no functional category survived correction for multiple testing, and consumption on the two sugar foods mapped to almost entirely separate sets of loci. Food intake therefore behaves as a classic polygenic trait, in which many variants of individually small effect contribute to variation in food intake. Whether the contributing genes act through diverse processes or through a shared one that current annotation does not capture cannot be determined here, given the number of uncharacterized genes within our gene pool.

Consumption of the two sugar foods was positively correlated across lines (r = 0.64 in males, 0.61 in females), so lines that ate more on one food tended to eat more on the other. The correlation explains only about 40% of the variance, however, and the two sugar foods recovered almost no shared loci in the association analysis. The correlation and the mapping point the same way: part of the variation is a general tendency to eat more or less, but much of it depends on which food the fly is on. This has a practical implication. Most studies of feeding measure intake on a single food, and our results suggest such designs will miss loci that act only under other dietary conditions. Describing the genetic architecture of feeding may therefore require sampling more than one food, even in a panel as well characterized as the DGRP.

### DGRP and GWAS

The DGRP has been used extensively to study the genetic basis of metabolic and behavioural traits (Francis et al., 2021; Garlapow et al., 2015; Jehrke et al., 2018; Unckless et al., 2015). To our knowledge, this is the first association study of food intake in the DGRP across two dietary sugar concentrations, and the first to map the compensatory adjustment between them. Measuring both sexes allowed sex-specific and diet-dependent effects to be distinguished. Consumption was heritable in both sexes and on both foods, and traits with heritability are those most likely to yield associated loci (Visscher et al., 2008; Zaitlen & Kraft, 2012), which justified the use of GWAS. Rather than analysing a single phenotype, we ran six analyses spanning sex, sugar foods and the low:high ratio, and we combined the resulting candidates into a single list.

This broadened the range of conditions sampled. Consumption on the two foods was only partly correlated, and the analyses shared almost no loci, so a single-diet study would likely have found a different set of genes.

We functionally validated 76 of the 118 candidate genes by RNAi knockdown. Knockdown of *CG42673* increased consumption in every combination of driver, sex, and sugar foods, and *mspo* and *Tmtc2* in most. *CG42673* and *Tmtc2* were retested in independent experiments. Under ubiquitous knockdown, the *CG42673* effect reproduced in both sexes, while the *Tmtc2* effect reproduced in females on the low-sugar food only and was absent in males in both experiments. The gene *mspo* has not yet been retested, and remains a strong candidate rather than a confirmed one. That knockdown raises intake indicates these genes are normally required to restrain consumption, but where in the process they act is not established. The same phenotype could arise from impaired satiety signalling, from altered nutrient absorption, or from a higher metabolic demand. Several candidates overlap with genes reported in earlier works. Both *mspo* and *Eip75B* were among the candidates of Garlapow et al. (2015) and Jehrke et al. (2018), and *Kirre* and *Sema*-*1a* have been implicated in more than one study (Francis et al., 2021; Garlapow et al., 2015; Jehrke et al., 2018). Recurrence across independent panels and phenotypes strengthens the case that these loci contribute to variation in feeding. It is important to note that the direction and magnitude of RNAi effects were not always consistent with those predicted by natural variants in the GWAS. Such discrepancies are expected, as RNAi knockdown globally reduces gene expression, whereas natural variants may alter traits through more subtle mechanisms such as regulatory changes or even increased expression (Garlapow et al., 2015). In addition, the genetic background of the RNAi lines differs from the DGRP, which may further contribute to variation in effect size or direction.

Two questions remain open. The first is where these genes are required. Ubiquitous knockdown reproducibly increased intake but pan-neuronal knockdowns gave inconsistent results across experiments. A second independent driver, *nSyb*-GAL4 produced no consistent effect across two independent trials, so the tissue of action is unresolved. The second is how these genes relate to established nutrient-sensing and neuroendocrine pathways. Answering both will be necessary to explain how natural variation at these loci translates into differences in how much a fly eats.

### Limitations

Several limitations should be noted. Functional testing covered 76 of the 118 candidate genes because 25 had no available RNAi line, and knockdown of 17 was lethal. So genes essential for viability could not be tested by this approach even if they contribute to variation in feeding. Each gene was targeted with a single RNAi line, without a second independent line or genetic rescue, so off-target effects cannot be formally excluded. Confirmatory experiments were limited to *CG42673* and *Tmtc2*. The gene *mspo*, which affected consumption in seven of eight conditions in the primary screen, has yet to be retested. The tissue in which the confirmed genes act is also unresolved, since ubiquitous and pan-neuronal knockdown gave different results. Finally, the DGRP comprises inbred, laboratory-adapted lines whose responses may not reflect those of outbred populations in the wild.

## Conclusions

Food consumption in *Drosophila* is shaped by diet, sex, and genetic background together. Flies compensate for dietary sugar by adjusting how much they eat, and the direction of this response proved to be very stable (it held across 14 days of exposure, in every wild-type strain, in 175 of 176 DGRP lines). However, its magnitude varied several-fold between genotypes, and that variation is heritable. Mapping this variation across the two sugar conditions identified 118 candidate genes. Consumption on low- and high-sugar food proved to be only partly shared genetically, with the two conditions recovering almost entirely separate loci, so dietary context shapes not only how much a fly eats but which genes contribute to that difference. Knockdown of *CG42673* increased consumption in every condition of the primary screen, and the effect was reproduced under ubiquitous knockdown in both sexes. *Tmtc2* knockdown reproduced in females on the low-sugar food. The gene *mspo* affected consumption in most conditions of the primary screen and remains a strong candidate. *CG42673* and *Tmtc2* are therefore normally required to limit food intake, although the tissue in which they act and the mechanism involved remain to be established. More broadly, these results show that mapping natural variation can identify feeding genes that no candidate-based approach would have tested. Understanding how genetic variation shapes food intake in flies may inform work on why humans differ in their response to sugar-rich foods.

## Acknowledgements

Stocks obtained from the Bloomington *Drosophila* Stock Center (NIH P40OD018537) were used in this study. The authors thank the University of Louisville’s Mentored Research Program and Summer Research Opportunity Program, as well as the Kentucky IDeA Network of Biomedical Research Excellence (KY INBRE) Summer Undergraduate Research Program, for supporting undergraduate research.

## Data availability

All fly stocks used in this study are available from the Bloomington *Drosophila* Stock Center, and stock numbers are listed in Table S3. DGRP genotypes and the association pipeline used here are publicly available at http://dgrp2.gnets.ncsu.edu. Phenotypic data generated in this study are available from the corresponding author on reasonable request.

## Author contributions

M.O. and D.-S.H. conceptualized and designed the study. M.O. performed the core experiments, analyzed the data, and drafted the manuscript. D.-S.H. supervised the project and revised the manuscript. L.F., S.A., M.W., S.A., and N.W. contributed to conducting the experiments. All authors approved the final version of the manuscript.

