## Supplementary figures and tables for "Novel Regulators of Sugar-dependent Food Consumption in *Drosophila melanogaster*"

**Figure S1. Food consumption of individual DGRP lines on low- and high-sugar food.**

Mean consumption of each DGRP line in males (a) and females (b), on food containing 5% (w/v) yeast and either 5% (low, blue) or 20% (w/v) sucrose (high, red). Each point is the mean of five biological replicates of 3–5 flies. Lines are shown in stock order rather than ranked. The same data ranked by low-sugar consumption appear in Figure 2a, b.

S1A

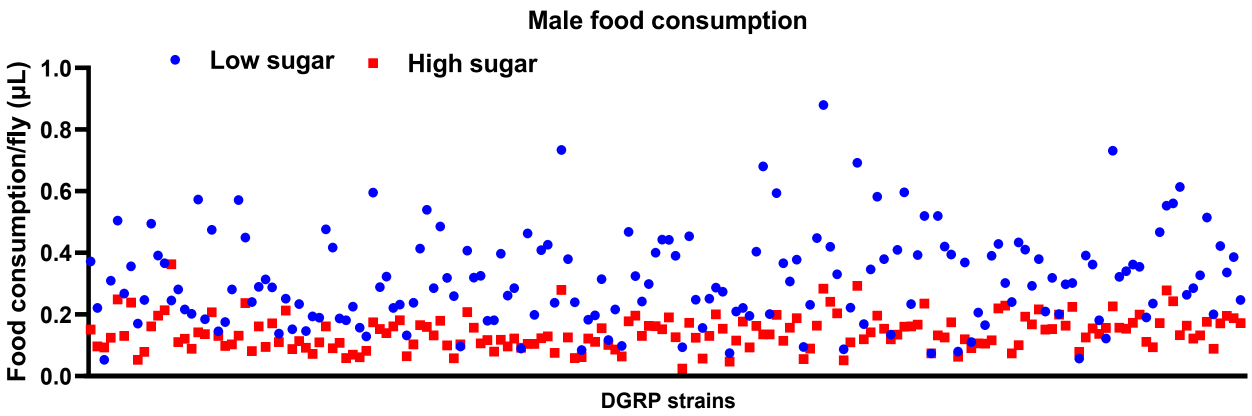


S1B

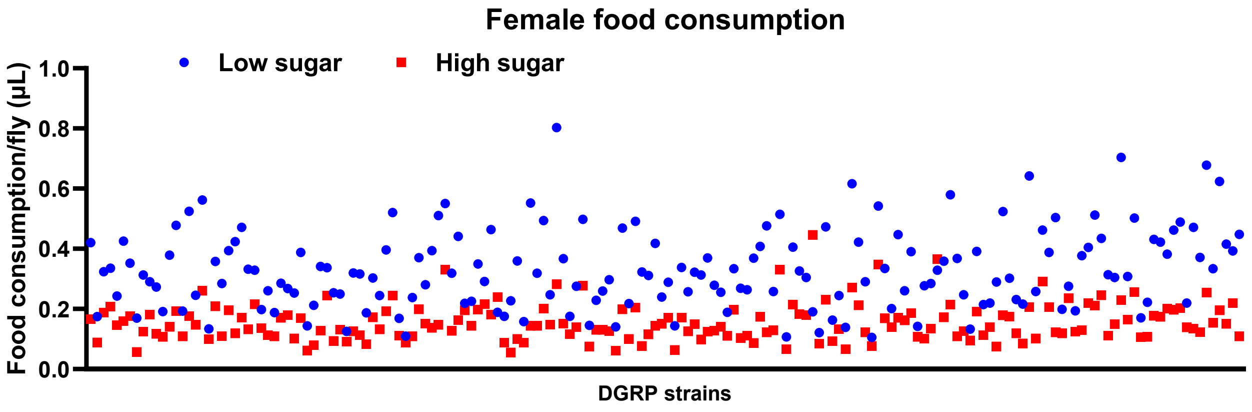

**Figure S2. Food consumption of the RNAi screen under ubiquitous knockdown.**

Absolute consumption of each RNAi line under arm-GAL4 in males (a) and females (b), on low-sugar (blue) and high-sugar (red) food. Bars show mean with individual replicates overlaid. Every line consumed more low- than high-sugar food in both sexes. Fold changes relative to the landing-site controls are in Figure 4a, b.

S2A

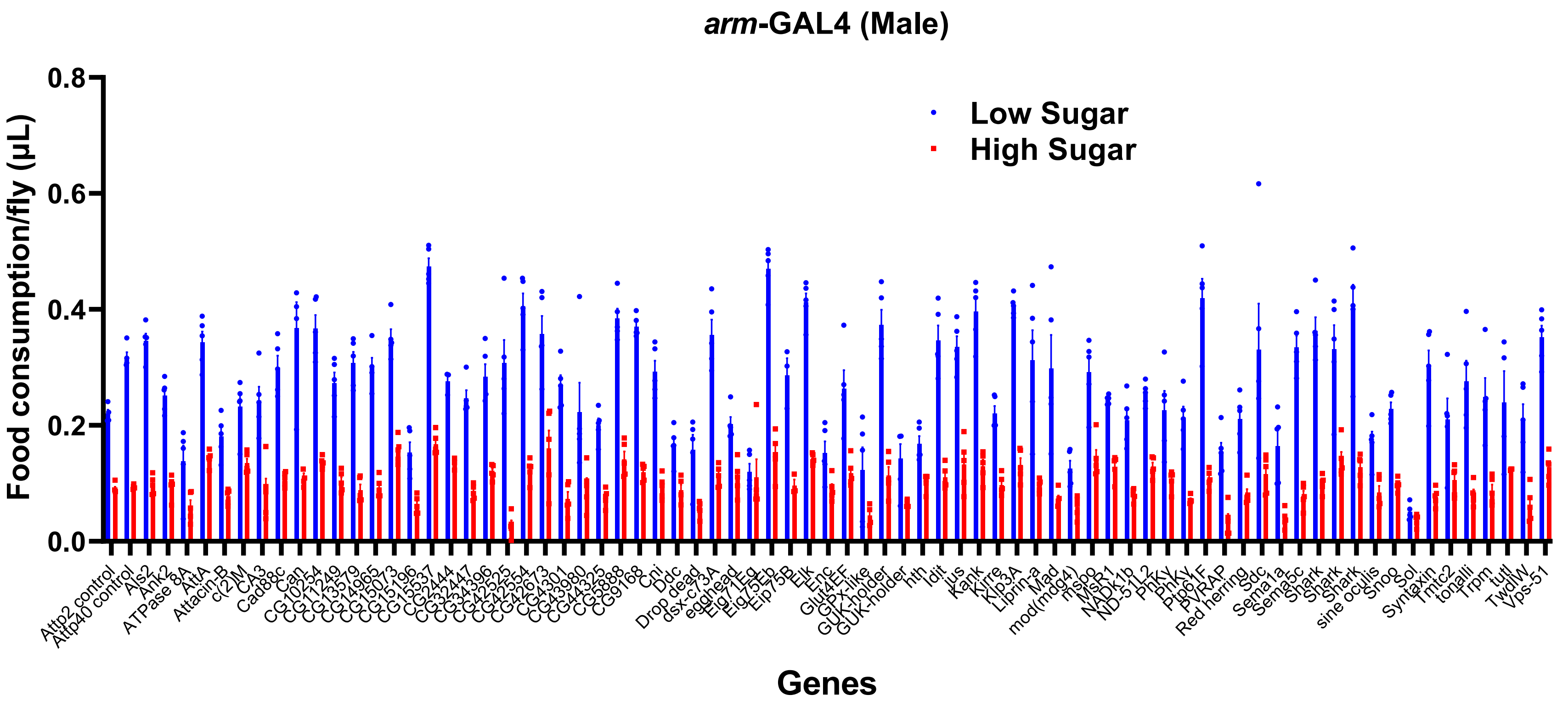

S2B

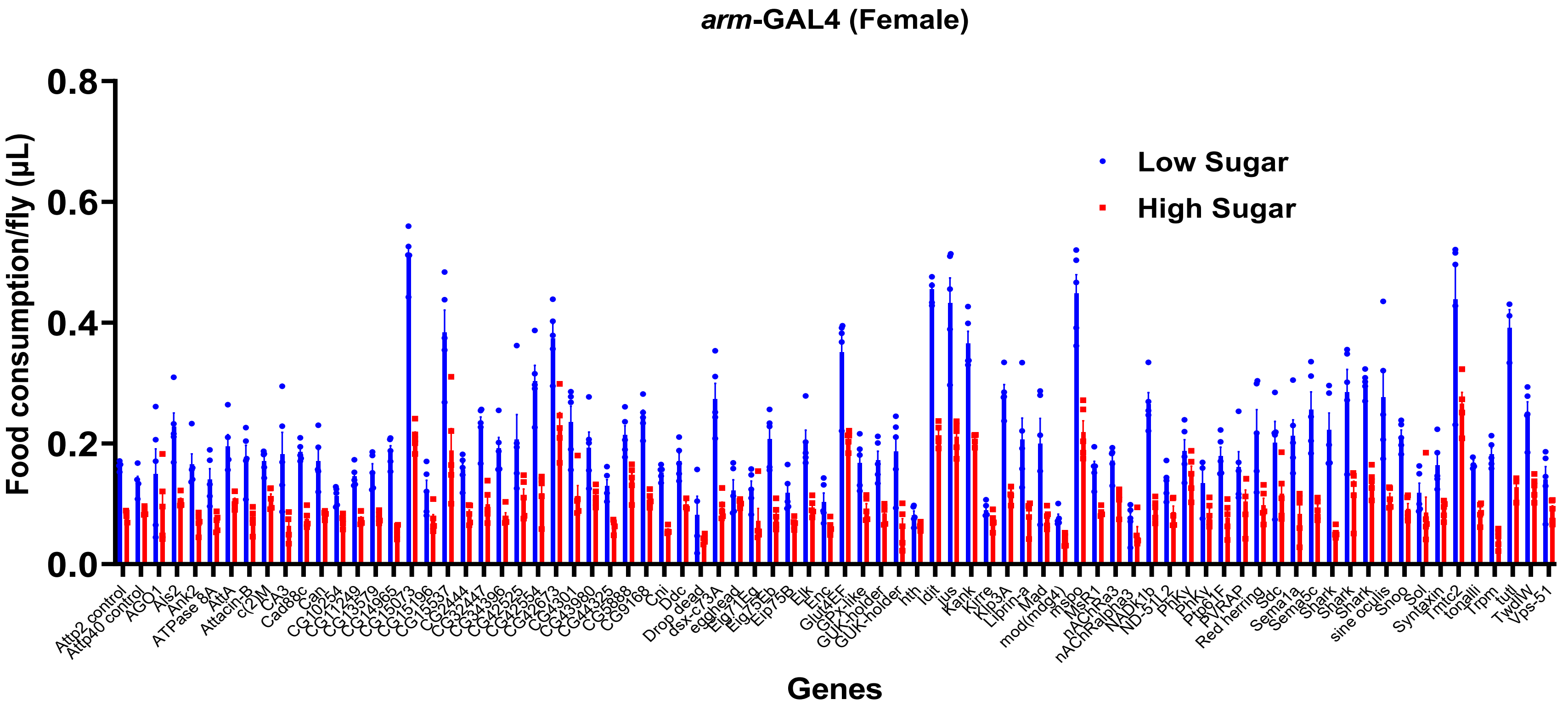

*arm*-GAL4 (Female)
Low Sugar
High Sugar
Food consumption/fly (µL)
0.8
0.6
0.4
0.2
0.0
Attp2 control
Attp40 control
AGO1
Als2
Ank2
ATPase 8A
AttA
Attacin-B
c(2)M
CA3
Cad88c
Can
CG10254
CG11249
CG13579
CG14965
CG15073
CG15196
CG15537
CG2444
CG32447
CG34396
CG42525
CG42554
CG42673
CG4301
CG43980
CG44325
CG5888
CG9168
Cni
Ddc
Drop dead
dsx-c73A
egghead
Eig71Eg
Eig75Eb
Eip75B
Elk
Enc
Glut4EF
GPx-like
GUK-holder
GUK-holder
hth
Idit
jus
Kank
Kirre
Klp3A
Liprin-a
Mad
mod(mdg4)
mspo
MsR1
nAChRa3
nAChRalpha3
NADk1b
ND-51L2
PhKy
PhKy
Ptp61F
PVRAP
Red herring
Sdc
Sema1a
Sema5c
Shark
Shark
Shark
sine oculis
Snoo
Sol
Syntaxin
Tmtc2
tonalli
Trpm
tutl
TwdlW
Vps-51
Genes

**Figure S3. Food consumption of the RNAi screen under pan-neuronal knockdown.**

Absolute consumption of each RNAi line under elav-GAL4 in males (a) and females (b), on low-sugar (blue) and high-sugar (red) food.

Bars show mean with individual replicates overlaid. Fold changes relative to the landing-site controls are in Figure 4c, d.

S3A

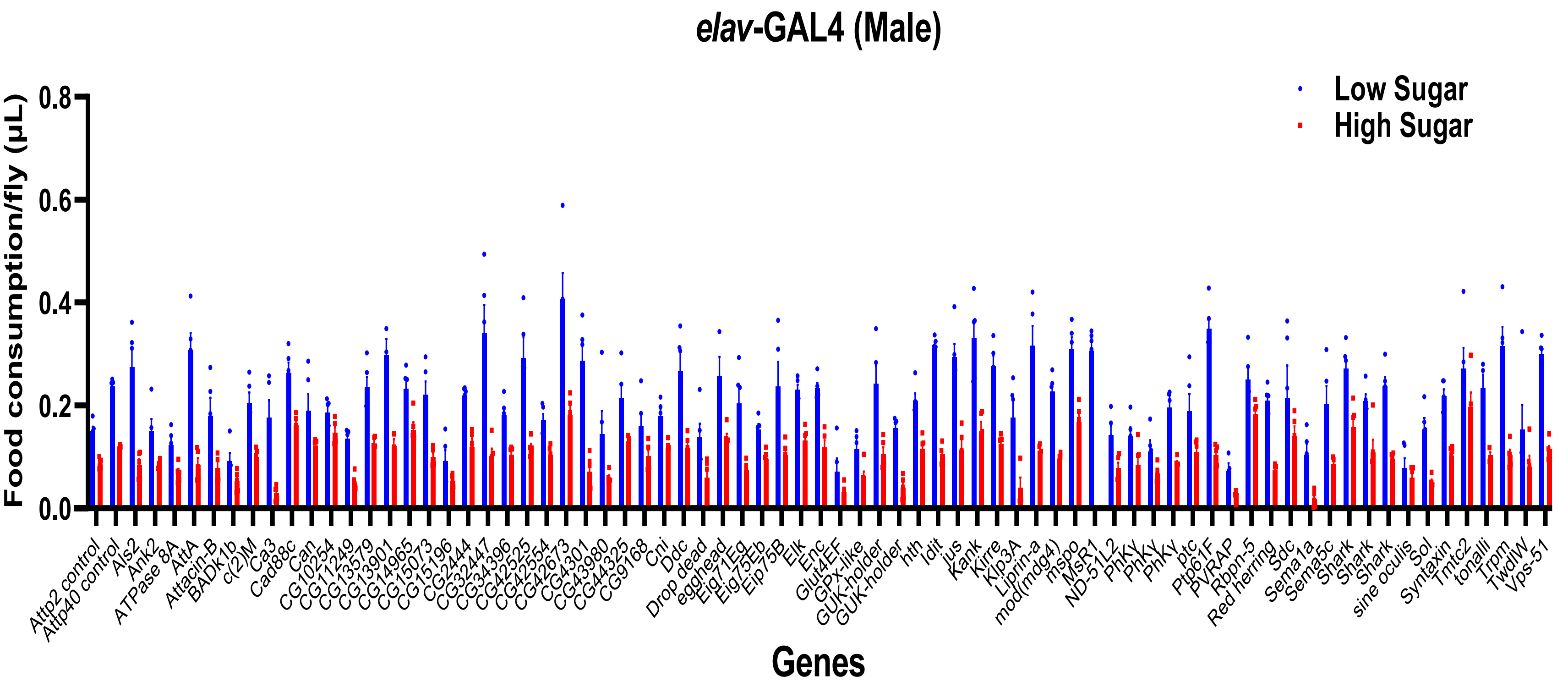

S3B

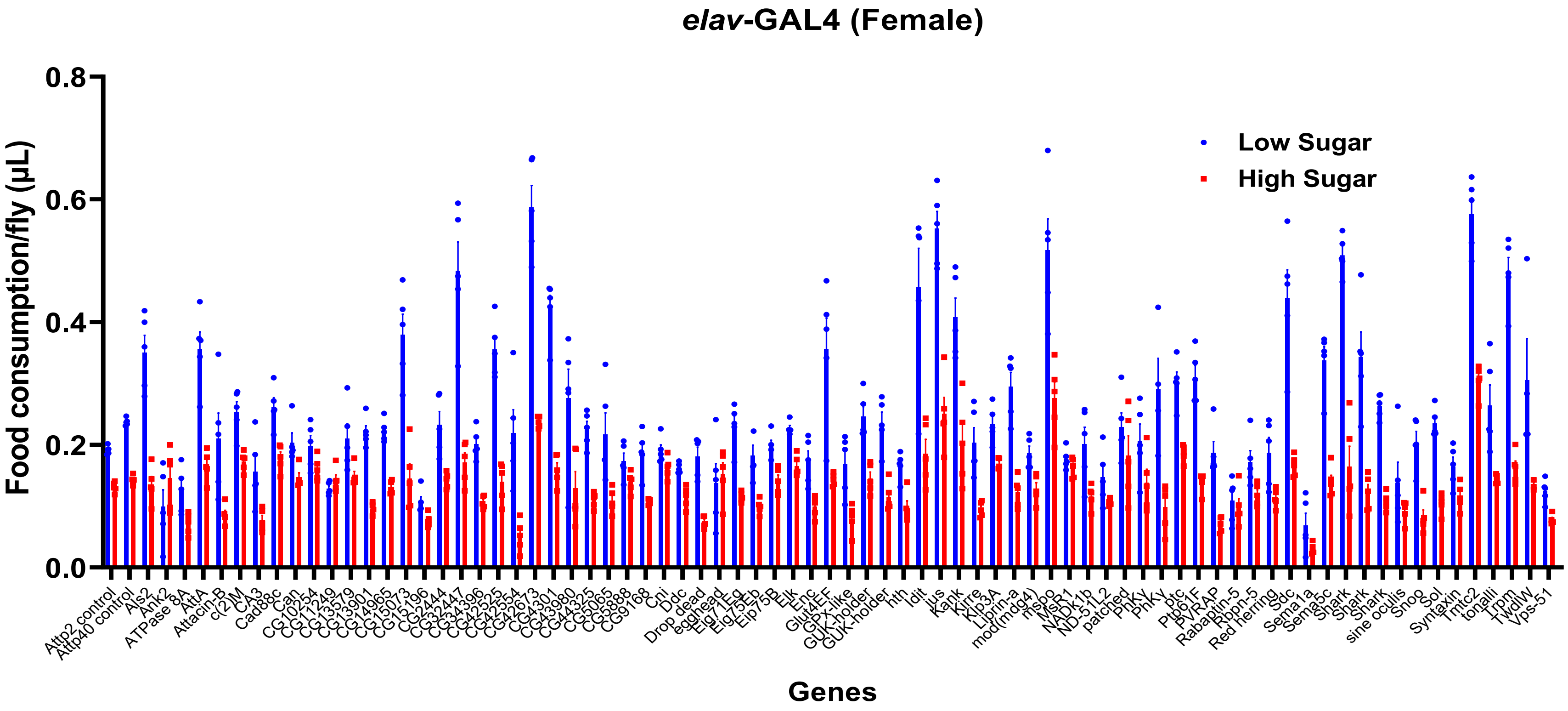

elav-GAL4 (Female)
Food consumption/fly (μL)
0.8
0.6
0.4
0.2
0.0
Low Sugar
High Sugar
Genes
Attp2 control
Attp40 control
Als2
Ank2
ATPase 8A
AttA
Attacin-B
c(2)M
CA3
Cad88c
Can
CG10254
CG11249
CG13579
CG13901
CG14965
CG15073
CG15196
CG2444
CG32447
CG34396
CG42525
CG42554
CG42673
CG4301
CG43980
CG44325
CG5065
CG5888
CG9168
Cni
Ddc
Drop dead
egghead
Eig71Eg
Eig75Eb
Eip75B
Elk
Enc
Glut4EF
GPx-like
GUK-holder
GUK-holder
hth
Idit
jus
Kank
Kirre
Klp3A
Liprin-a
mod(mdg4)
mspo
MsR1
NADK1b
ND-51L2
patched
PhKγ
PhKγ
ptc
Ptp61F
PVRAP
Rabaptin-5
Rbpn-5
Red herring
Sdc
Sema1a
Sema5c
Shark
Shark
Shark
sine oculis
Snoo
Sol
Syntaxin
Tmtc2
tonalli
Trpm
Twdlw
Vps-51

**Figure S4. Knockdown of CG42673 and Tmtc2 under additional neuronal and glial drivers.**

Food consumption in females following knockdown with elav-GAL4 in two further independent experiments (a, b), nSyb-GAL4 (BDSC 51941) in two independent trials (c, d), and repo-GAL4 (e). Bars show mean ± SEM with individual replicates overlaid, n = 5 per genotype and diet.

S4A

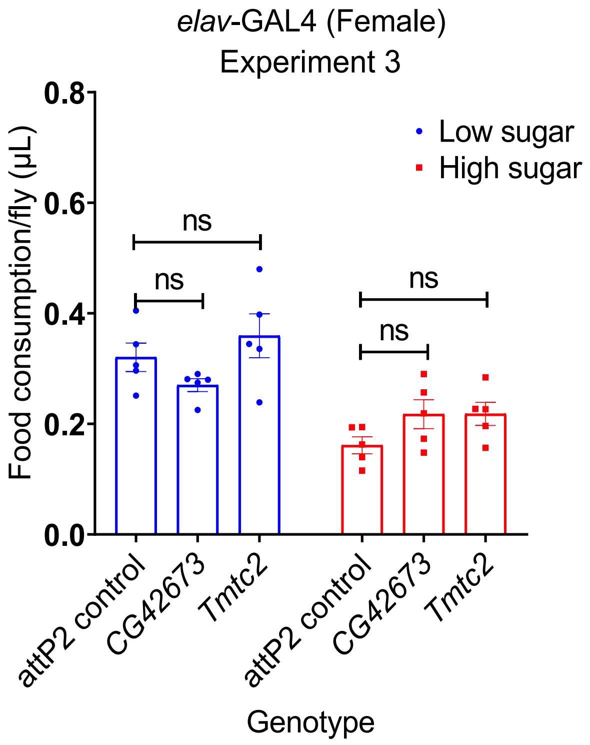


S4B

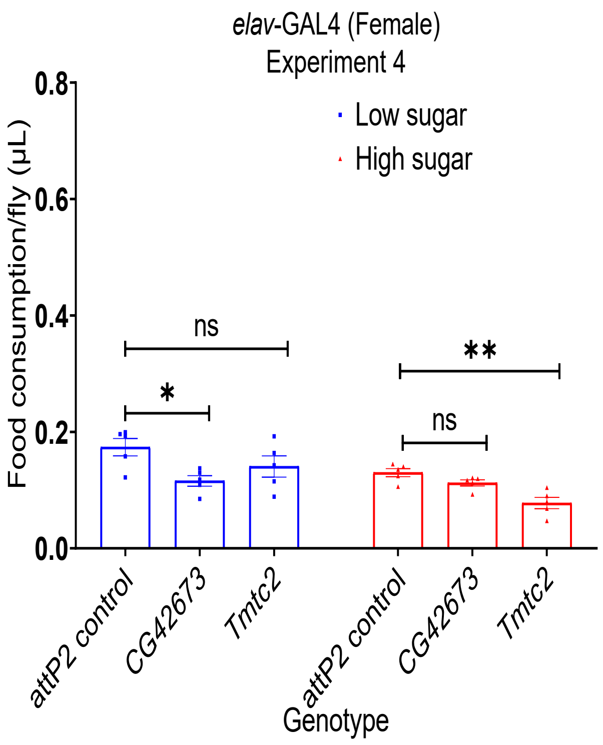

S4C

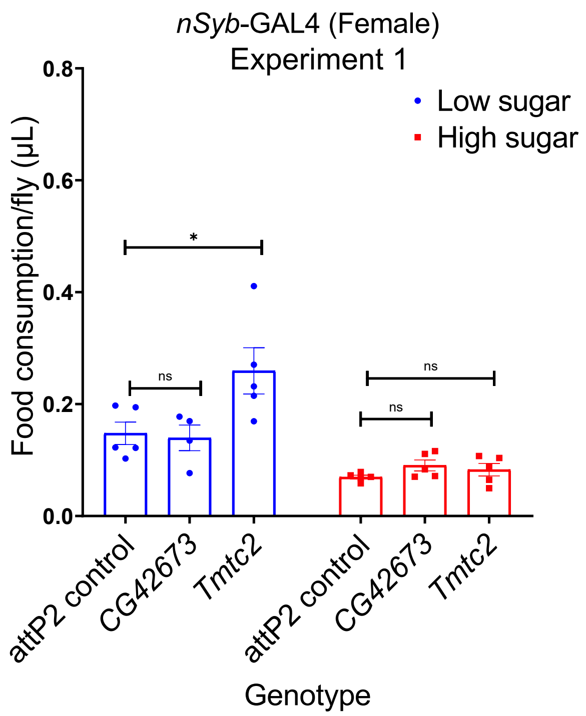

nSyb-GAL4 (Female)
Experiment 1
Low sugar
High sugar
Food consumption/fly (μL)
0.8
0.6
0.4
0.2
0.0
*
ns
ns
ns
attP2 control
CG42673
Tmtc2
attP2 control
CG42673
Tmtc2
Genotype


S4D

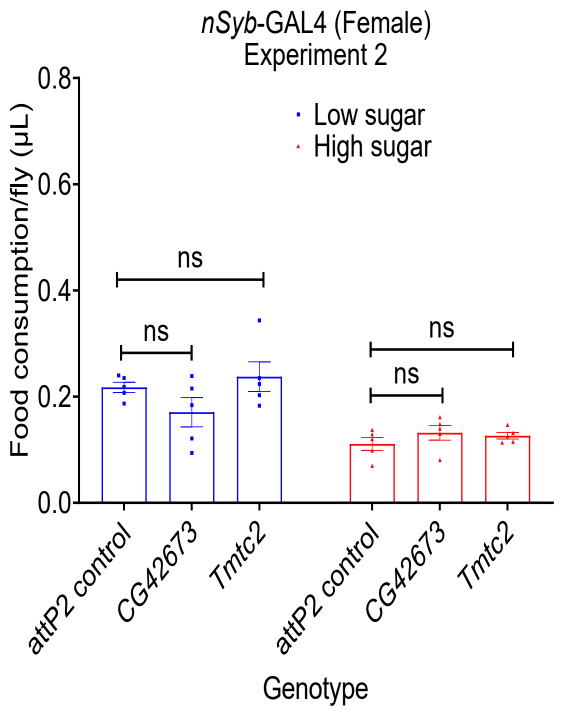

nSyb-GAL4 (Female)
Experiment 2
Low sugar
High sugar
Food consumption/fly (μL)
0.8
0.6
0.4
0.2
0.0
ns
ns
ns
ns
attP2 control
CG42673
Tmtc2
attP2 control
CG42673
Tmtc2
Genotype


S4E

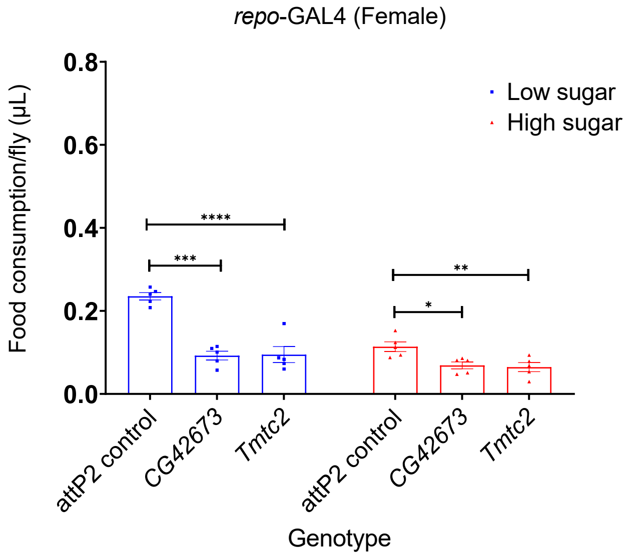

repo-GAL4 (Female)
Low sugar
High sugar
Food consumption/fly (μL)
0.8
0.6
0.4
0.2
0.0
****
***
**
*
attP2 control
CG42673
Tmtc2
attP2 control
CG42673
Tmtc2
Genotype

**Table S1. Covariate effects on food consumption phenotypes prior to association analysis.**

P values from Type III ANOVA testing the effect of *Wolbachia* infection status and five common polymorphic inversions on each phenotype. All phenotypes were adjusted for the full covariate set before association testing, irrespective of significance.

| **Phenotype** | ***Wolbachia*** | ***In(2L)t*** | ***In(2R)NS*** | ***In(3R)P*** | ***In(3R)K*** | ***In(3R)Mo*** |
|---|---|---|---|---|---|---|
| Female, low sugar | 0.339 | 0.358 | 0.122 | 0.838 | 0.630 | 0.573 |
| Female, high sugar | 0.934 | 0.923 | 0.235 | 0.791 | 0.660 | 0.823 |
| Female, low:high ratio | 0.152 | 0.253 | 0.053 | 0.946 | 0.728 | 0.291 |
| Male, low sugar | 0.166 | 0.073 | **0.039** | 0.225 | 0.607 | 0.295 |
| Male, high sugar | 0.297 | 0.152 | 0.115 | 0.784 | 0.977 | 0.720 |
| Male, low:high ratio | **0.018** | 0.502 | 0.064 | 0.134 | 0.799 | 0.317 |

Bold indicates $P < 0.05$.

**Table S2. Functional annotation enrichment among the 118 candidate genes.**

All annotation terms reaching nominal P < 0.05 in DAVID, tested against the full annotated Drosophila melanogaster genome as background. Of 118 genes submitted, 96 were annotated for biological process, 101 for cellular component and 85 for molecular function. Rows in bold with grey shading are significant after Benjamini–Hochberg correction (adjusted P < 0.05); all other rows are nominally enriched only. Count, genes from the submitted list carrying the term; List total, annotated genes in the submitted list for that category; Pop hits, genes carrying the term in the background; Pop total, annotated background genes for that category.

| Category | Term | Count | List total | Pop hits | Pop total | P value | Benjamini | Fold enrichment |
|---|---|---|---|---|---|---|---|---|
| **UP_SEQ_FEATURE** | **COMPBIAS: Polar residues** | **50** | **96** | **3379** | **11956** | **1.29E-06** | **2.60E-04** | **1.84** |
| **INTERPRO** | **Insect_Unk** | **5** | **100** | **11** | **11761** | **1.49E-06** | **5.44E-04** | **53.46** |
| **UP_SEQ_FEATURE** | **REGION: Disordered** | **79** | **96** | **7203** | **11956** | **3.67E-06** | **3.71E-04** | **1.37** |
| **UP_SEQ_FEATURE** | **COMPBIAS: Low complexity** | **50** | **96** | **3557** | **11956** | **6.41E-06** | **4.31E-04** | **1.75** |
| **GOTERM_BP_DIRECT** | **defense response to bacterium** | **7** | **96** | **79** | **12487** | **2.94E-05** | **1.32E-02** | **11.53** |
| GOTERM_CC_DIRECT | presynaptic membrane | 4 | 101 | 38 | 12854 | 3.16E-03 | 4.08E-01 | 13.40 |
| UP_KW_MOLECULAR_FUNCTION | Developmental protein | 13 | 45 | 687 | 5715 | 4.54E-03 | 8.63E-02 | 2.40 |
| GOTERM_BP_DIRECT | positive regulation of synaptic assembly at neuromuscular junction | 4 | 96 | 45 | 12487 | 4.80E-03 | 1.00E+00 | 11.56 |
| GOTERM_BP_DIRECT | phospholipid transport | 3 | 96 | 21 | 12487 | 1.09E-02 | 1.00E+00 | 18.58 |
| GOTERM_BP_DIRECT | axon guidance | 6 | 96 | 176 | 12487 | 1.10E-02 | 1.00E+00 | 4.43 |

| Category | Term | Count | List total | Pop hits | Pop total | P value | Benjamini | Fold enrichment |
|---|---|---|---|---|---|---|---|---|
| UP_KW_DOMAIN | ANK repeat | 4 | 74 | 52 | 8084 | 1.14E-02 | 1.71E-01 | 8.40 |
| GOTERM_BP_DIRECT | eye development | 3 | 96 | 26 | 12487 | 1.65E-02 | 1.00E+00 | 15.01 |
| GOTERM_BP_DIRECT | regulation of membrane potential | 3 | 96 | 27 | 12487 | 1.78E-02 | 1.00E+00 | 14.45 |
| GOTERM_BP_DIRECT | follicle cell of egg chamber development | 4 | 96 | 78 | 12487 | 2.15E-02 | 1.00E+00 | 6.67 |
| COG_ONTOLOGY | Cell division and chromosome partitioning | 3 | 8 | 44 | 1239 | 2.31E-02 | 9.25E-02 | 10.56 |
| UP_SEQ_FEATURE | DOMAIN: Attacin N-terminal | 2 | 96 | 3 | 11956 | 2.37E-02 | 7.96E-01 | 83.03 |
| GOTERM_MF_DIRECT | co-SMAD binding | 2 | 85 | 4 | 12350 | 2.69E-02 | 1.00E+00 | 72.65 |
| GOTERM_BP_DIRECT | synaptic target recognition | 3 | 96 | 34 | 12487 | 2.74E-02 | 1.00E+00 | 11.48 |
| UP_SEQ_FEATURE | COMPBIAS: Basic residues | 22 | 96 | 1726 | 11956 | 2.82E-02 | 7.96E-01 | 1.59 |
| UP_SEQ_FEATURE | TOPO_DOM: Cytoplasmic | 10 | 96 | 544 | 11956 | 2.87E-02 | 7.96E-01 | 2.29 |
| GOTERM_BP_DIRECT | insect visual primordium formation | 2 | 96 | 4 | 12487 | 3.01E-02 | 1.00E+00 | 65.04 |
| UP_SEQ_FEATURE | DOMAIN: Attacin C-terminal | 2 | 96 | 4 | 11956 | 3.14E-02 | 7.96E-01 | 62.27 |
| UP_SEQ_FEATURE | COMPBIAS: Gly residues | 14 | 96 | 928 | 11956 | 3.15E-02 | 7.96E-01 | 1.88 |
| INTERPRO | Attacin_N | 2 | 100 | 4 | 11761 | 3.33E-02 | 1.00E+00 | 58.80 |
| GOTERM_BP_DIRECT | compound eye morphogenesis | 4 | 96 | 93 | 12487 | 3.39E-02 | 1.00E+00 | 5.59 |
| INTERPRO | Ankyrin_rpt | 4 | 100 | 85 | 11761 | 3.48E-02 | 1.00E+00 | 5.53 |
| GOTERM_CC_DIRECT | transcription regulator complex | 4 | 101 | 92 | 12854 | 3.49E-02 | 1.00E+00 | 5.53 |
| INTERPRO | Ankyrin_rpt-contain_sf | 4 | 100 | 86 | 11761 | 3.58E-02 | 1.00E+00 | 5.47 |
| UP_SEQ_FEATURE | COMPBIAS: Acidic residues | 19 | 96 | 1454 | 11956 | 3.62E-02 | 8.12E-01 | 1.63 |
| GOTERM_CC_DIRECT | neuromuscular junction | 4 | 101 | 95 | 12854 | 3.78E-02 | 1.00E+00 | 5.36 |

| Category | Term | Count | List total | Pop hits | Pop total | P value | Benjamini | Fold enrichment |
|---|---|---|---|---|---|---|---|---|
| GOTERM_MF_DIRECT | ATPase-coupled intramembrane lipid carrier activity | 2 | 85 | 6 | 12350 | 4.01E-02 | 1.00E+00 | 48.43 |
| GOTERM_CC_DIRECT | membrane | 25 | 101 | 2162 | 12854 | 4.09E-02 | 1.00E+00 | 1.47 |
| GOTERM_CC_DIRECT | plasma membrane | 21 | 101 | 1722 | 12854 | 4.13E-02 | 1.00E+00 | 1.55 |
| INTERPRO | Attacin_C | 2 | 100 | 5 | 11761 | 4.14E-02 | 1.00E+00 | 47.04 |
| SMART | ANK | 4 | 52 | 82 | 5399 | 4.17E-02 | 1.00E+00 | 5.06 |
| GOTERM_BP_DIRECT | segment polarity determination | 3 | 96 | 45 | 12487 | 4.59E-02 | 1.00E+00 | 8.67 |
| UP_SEQ_FEATURE | DOMAIN: P-type ATPase C-terminal | 2 | 96 | 6 | 11956 | 4.67E-02 | 8.58E-01 | 41.51 |
| UP_SEQ_FEATURE | DOMAIN: P-type ATPase N-terminal | 2 | 96 | 6 | 11956 | 4.67E-02 | 8.58E-01 | 41.51 |
| INTERPRO | P_typ_ATPase_c | 2 | 100 | 6 | 11761 | 4.95E-02 | 1.00E+00 | 39.20 |
| INTERPRO | P-type_ATPase_IV | 2 | 100 | 6 | 11761 | 4.95E-02 | 1.00E+00 | 39.20 |
| INTERPRO | P-type_ATPase_N | 2 | 100 | 6 | 11761 | 4.95E-02 | 1.00E+00 | 39.20 |

The seven genes annotated with defense response to bacterium (GO:0042742) are *Attacin-A* (FBgn0012042), *Attacin-B* (FBgn0041581), *Eig71Ea* (FBgn0004588), *Eig71Eb* (FBgn0004589), *Eig71Ef* (FBgn0004593), *Eig71Eg* (FBgn0004594) and *Eig71Eh* (FBgn0014848). *Attacin-A* and *Attacin-B* are tagged by a single shared variant (2R_10636472_SNP), and the five *Eig71E* genes lie in a single tandem cluster; this enrichment therefore reflects physical clustering of antimicrobial peptide genes rather than independent association at seven loci.

### Table S3. *Drosophila* stocks used in this study.

All stocks were obtained from the Bloomington Drosophila Stock Center (BDSC). Genotypes are given as listed by BDSC at the ti me of ordering.

#### (a) DGRP lines (n = 176)

| BDSC | Line | BDSC | Line | BDSC | Line | BDSC | Line |
|---|---|---|---|---|---|---|---|
| 25174 | DGRP-208 | 28182 | DGRP-370 | 29660 | DGRP-530 | 28227 | DGRP-761 |
| 25176 | DGRP-303 | 28183 | DGRP-371 | 37525 | DGRP-306 | 28229 | DGRP-776 |
| 25180 | DGRP-313 | 28185 | DGRP-374 | 55014 | DGRP-31 | 28231 | DGRP-787 |
| 25184 | DGRP-357 | 28186 | DGRP-377 | 55015 | DGRP-32 | 28239 | DGRP-810 |
| 25186 | DGRP-360 | 28189 | DGRP-382 | 55016 | DGRP-48 | 28240 | DGRP-812 |
| 25188 | DGRP-375 | 28191 | DGRP-385 | 55018 | DGRP-319 | 28246 | DGRP-837 |
| 25189 | DGRP-379 | 28194 | DGRP-392 | 55019 | DGRP-348 | 28250 | DGRP-853 |
| 25190 | DGRP-380 | 28196 | DGRP-426 | 55022 | DGRP-395 | 28251 | DGRP-855 |
| 25191 | DGRP-391 | 28197 | DGRP-440 | 55023 | DGRP-397 | 28252 | DGRP-857 |
| 25194 | DGRP-437 | 28198 | DGRP-441 | 55027 | DGRP-559 | 28254 | DGRP-879 |
| 25198 | DGRP-555 | 28199 | DGRP-443 | 55030 | DGRP-627 | 28255 | DGRP-882 |
| 25200 | DGRP-707 | 28202 | DGRP-491 | 55031 | DGRP-630 | 28260 | DGRP-897 |
| 25203 | DGRP-732 | 28204 | DGRP-502 | 55032 | DGRP-634 | 28261 | DGRP-900 |
| 25204 | DGRP-765 | 28206 | DGRP-509 | 83728 | DGRP-352 | 29654 | DGRP-320 |
| 25205 | DGRP-774 | 28207 | DGRP-531 | 83729 | DGRP-737 | 29655 | DGRP-321 |
| 25208 | DGRP-820 | 28208 | DGRP-535 | 25177 | DGRP-304 | 29656 | DGRP-405 |
| 25209 | DGRP-852 | 28212 | DGRP-584 | 25179 | DGRP-307 | 55017 | DGRP-100 |
| 25210 | DGRP-859 | 28215 | DGRP-595 | 25181 | DGRP-315 | 55020 | DGRP-354 |
| 25744 | DGRP-705 | 28218 | DGRP-703 | 25182 | DGRP-324 | 55021 | DGRP-390 |
| 28122 | DGRP-21 | 28223 | DGRP-738 | 25185 | DGRP-358 | 55024 | DGRP-505 |
| 28126 | DGRP-41 | 28230 | DGRP-783 | 25192 | DGRP-399 | 55026 | DGRP-551 |
| 28128 | DGRP-45 | 28232 | DGRP-790 | 25193 | DGRP-427 | 55028 | DGRP-566 |
| 28132 | DGRP-75 | 28233 | DGRP-796 | 25197 | DGRP-517 | 55038 | DGRP-355 |
| 28136 | DGRP-91 | 28234 | DGRP-801 | 25199 | DGRP-639 | 25183 | DGRP-335 |
| 28139 | DGRP-105 | 28235 | DGRP-802 | 25201 | DGRP-712 | 25187 | DGRP-362 |
| 28140 | DGRP-109 | 28236 | DGRP-804 | 25206 | DGRP-786 | 25195 | DGRP-486 |
| 28141 | DGRP-129 | 28237 | DGRP-805 | 28127 | DGRP-42 | 25207 | DGRP-799 |
| 28142 | DGRP-136 | 28238 | DGRP-808 | 28137 | DGRP-93 | 25445 | DGRP-365 |
| 28144 | DGRP-142 | 28241 | DGRP-818 | 28138 | DGRP-101 | 25745 | DGRP-714 |
| 28145 | DGRP-149 | 28242 | DGRP-819 | 28150 | DGRP-177 | 28123 | DGRP-26 |
| 28148 | DGRP-161 | 28243 | DGRP-821 | 28151 | DGRP-181 | 28135 | DGRP-88 |
| 28153 | DGRP-195 | 28244 | DGRP-822 | 28154 | DGRP-217 | 28143 | DGRP-138 |
| 28156 | DGRP-227 | 28249 | DGRP-850 | 28161 | DGRP-239 | 28146 | DGRP-153 |
| 28157 | DGRP-228 | 28256 | DGRP-884 | 28164 | DGRP-280 | 28152 | DGRP-189 |
| 28165 | DGRP-287 | 28257 | DGRP-890 | 28176 | DGRP-350 | 28219 | DGRP-716 |

| | | | | | | | |
|---|---|---|---|---|---|---|---|
| 28166 | DGRP-309 | 28262 | DGRP-907 | 28184 | DGRP-373 | 28247 | DGRP-843 |
| 28167 | DGRP-317 | 28263 | DGRP-908 | 28190 | DGRP-383 | 28253 | DGRP-861 |
| 28168 | DGRP-318 | 28265 | DGRP-913 | 28192 | DGRP-386 | 28258 | DGRP-892 |
| 28171 | DGRP-332 | 28275 | DGRP-235 | 28203 | DGRP-492 | 28264 | DGRP-911 |
| 28172 | DGRP-336 | 28276 | DGRP-310 | 28211 | DGRP-563 | 28274 | DGRP-85 |
| 28173 | DGRP-338 | 28278 | DGRP-409 | 28213 | DGRP-589 | 29657 | DGRP-406 |
| 28174 | DGRP-340 | 29651 | DGRP-40 | 28220 | DGRP-721 | 29658 | DGRP-439 |
| 28179 | DGRP-359 | 29652 | DGRP-57 | 28224 | DGRP-748 | 55025 | DGRP-528 |
| 28180 | DGRP-361 | 29659 | DGRP-513 | 28226 | DGRP-757 | 28129 | DGRP-59 |

**(b) Wild-type strains (n = 8)**

| BDSC stock | Strain | Note |
|---|---|---|
| 5905 | w1118 | Used in Figure 1a-c |
| 5 | Oregon-R-C | |
| 4265 | Amherst_3 | |
| 4268 | Lausanne-S | |
| 4270 | Samarkand | |
| 4271 | Swedish-C | |
| 4272 | Urbana-S | |
| 60741 | Zimbabwe-S-29 | |

**(c) GAL4 drivers (n = 5)**

| BDSC stock | Driver | Genotype |
|---|---|---|
| 4414 | *act*-GAL4 | y[1] w[*]; P{w[+mC]=Act5C-GAL4}25FO1/CyO, y[+] |
| 1560 | *arm*-GAL4 | w[*]; P{w[+mW.hs]=GAL4-arm.S}11 |
| 458 | *elav*-GAL4 | P{w[+mW.hs]=GawB}elav[C155] |
| 51941 | *nSyb*-GAL4 | y[1] w[1118]; P{y[+t7.7] w[+mC]=nSyb-GAL4.P}attP2 |
| 7415 | *repo*-GAL4 | w[1118]; P{w[+m*]=GAL4}repo/TM3, Sb[1] |

**(d) Landing-site controls (n = 2)**

| BDSC stock | Control | Genotype |
|---|---|---|
| 36303 | attP2 control | y[1] v[1]; P{y[+t7.7]=CaryP}attP2 |
| 36304 | attP40 control | y[1] v[1]; P{y[+t7.7]=CaryP}attP40 |

#### (e) UAS-RNAi lines (n = 79)

| Target gene | BDSC stock | Landing site | Genotype |
|---|---|---|---|
| *AGO1* | 53293 | attP40 | y[1] sc[*] v[1] sev[21]; P{TRiP.HMS00610}attP2 |
| *Als2* | 28533 | attP2 | y[1] v[1]; P{TRiP.HM05019}attP2 |
| *Ank2* | 29438 | attP2 | y[1] v[1]; P{y[+t7.7] v[+t1.8]=TRiP.JF03374}attP2 |
| *ATPase 8A* | 28558 | attP2 | y[1] v[1]; P{TRiP.HM05044}attP2/TM3, Sb[1] |
| *AttA* | 56904 | attP40 | y[1] sc[*] v[1] sev[21]; P{y[+t7.7] v[+t1.8]=TRiP.HMC04126}attP40 |
| *AttB* | 57392 | attP40 | y[1] sc[*] v[1] sev[21]; P{TRiP.HMC04696}attP40 |
| *c(2)M* | 43977 | attP2 | y[1] v[1]; P{y[+t7.7] v[+t1.8]=TRiP.GL01587}attP2/TM6C, Sb[1] Tb[1] |
| *CA3* | 66985 | attP40 | y[1] sc[*] v[1] sev[21]; P{TRiP.HMS05451}attP40 |
| *Cad88c* | 29303 | attP2 | y[1] v[1]; P{y[+t7.7] v[+t1.8]=TRiP.JF02443}attP2 |
| *Can* | 67756 | attP40 | y[1] sc[*] v[1] sev[21]; P{y[+t7.7] v[+t1.8]=TRiP.HMS05778}attP40/CyO |
| *CG10254* | 51687 | attP40 | y[1] v[1]; P{y[+t7.7] v[+t1.8]=TRiP.GLC01787}attP40 |
| *CG11249* | 57240 | attP40 | y[1] sc[*] v[1] sev[21]; P{TRiP.HMC04626}attP40 |
| *CG13579* | 28644 | attP2 | y[1] v[1]; P{TRiP.JF03059}attP2 |
| *CG13901* | 66336 | attP2 | y[1] sc[*] v[1] sev[21]; P{y[+t7.7] v[+t1.8]=TRiP.HMC06270}attP2 |
| *CG14965* | 56871 | attP40 | y[1] v[1]; P{y[+t7.7] v[+t1.8]=TRiP.HMC03727}attP40 |
| *CG15073* | 29366 | attP2 | y[1] v[1]; P{TRiP.JF02530}attP2 |
| *CG15196* | 53688 | attP40 | y[1] v[1]; P{TRiP.HMJ21648}attP40/CyO |
| *CG2444* | 58108 | attP40 | y[1] v[1]; P{y[+t7.7] v[+t1.8]=TRiP.HMJ22044}attP40 |
| *CG32447* | 60466 | attP40 | y[1] v[1]; P{TRiP.HMJ22835}attP40 |
| *CG34396* | 26011 | attP2 | y[1] v[1]; P{y[+t7.7] v[+t1.8]=TRiP.JF02036}attP2 |
| *CG42525* | 67375 | attP40 | y[1] sc[*] v[1] sev[21]; P{TRiP.HMC06483}attP40 |
| *CG42554* | 77451 | attP40 | y[1] sc[*] v[1] sev[21]; P{TRiP.HMC06583}attP40 |
| *CG42673* | 36629 | attP2 | y[1] sc[*] v[1] sev[21]; P{TRiP.GL00589}attP2 |
| *CG4301* | 64610 | attP40 | y[1] sc[*] v[1] sev[21]; P{TRiP.HMC05645}attP40 |
| *CG43980* | 57167 | attP40 | y[1] sc[*] v[1] sev[21]; P{TRiP.HMC04548}attP40 |
| *CG44325* | 43209 | attP40 | y[1] sc[*] v[1] sev[21]; P{TRiP.GL01554}attP40 |
| *CG5065* | 603574 | attP40 | y[1] v[1]; P{y[+t7.7] v[+t1.8]=TRiP.HMS06233}attP40/CyO |
| *CG5888* | 62175 | attP40 | y[1] sc[*] v[1] sev[21]; P{y[+t7.7] v[+t1.8]=TRiP.HMC05182}attP40/CyO |
| *CG9168* | 62976 | attP40 | y[1] v[1]; P{y[+t7.7] v[+t1.8]=TRiP.HMJ30053}attP40 |
| *Cni* | 38919 | attP2 | y[1] sc[*] v[1] sev[21]; P{y[+t7.7] v[+t1.8]=TRiP.GL00691}attP2 |
| *Ddc* | 51462 | attP40 | y[1] v[1]; P{y[+t7.7] v[+t1.8]=TRiP.HMC03200}attP40 |
| *Drop dead* | 66957 | attP40 | y[1] sc[*] v[1] sev[21]; P{TRiP.HMS05423}attP40 |
| *dsx-c73A* | 56987 | attP40 | y[1] sc[*] v[1] sev[21]; P{TRiP.HMC04427}attP40 |
| *egghead* | 28687 | attP2 | y[1] v[1]; P{TRiP.JF03102}attP2 |
| *Eig71Eb* | 56951 | attP40 | y[1] sc[*] v[1] sev[21]; P{y[+t7.7] v[+t1.8]=TRiP.HMC04391}attP40 |
| *Eig71Eg* | 56953 | attP40 | y[1] sc[*] v[1] sev[21]; P{TRiP.HMC04393}attP40 |
| *Eip75B* | 35780 | attP2 | y[1] sc[*] v[1] sev[21]; P{TRiP.GLC01418}attP2 |
| *Elk* | 25821 | attP2 | y[1] v[1]; P{y[+t7.7] v[+t1.8]=TRiP.JF01839}attP2 |

| | | | |
|---|---|---|---|
| *Enc* | 42797 | attP2 | y[1] v[1]; P{y[+t7.7] v[+t1.8]=TRiP.GL01168}attP2 |
| *Glut4EF* | 57461 | attP40 | y[1] sc[*] v[1] sev[21]; P{TRiP.HMC04771}attP40 |
| *Gpxl* | 56992 | attP40 | y[1] sc[*] v[1] sev[21]; P{TRiP.HMC04434}attP40 |
| *GUK-holder* | 42486 | attP40 | y[1] sc[*] v[1] sev[21]; P{TRiP.HMC03681}attP2 |
| *GUK-holder* | 55858 | attP2 | y[1] sc[*] v[1] sev[21]; P{TRiP.HMC03681}attP2 |
| *hth* | 34637 | attP2 | y[1] sc[*] v[1] sev[21]; P{TRiP.HMS01112}attP2 |
| *Idit* | 28823 | attP2 | y[1] v[1]; P{TRiP.JF02930}attP2 |
| *jus* | 28764 | attP2 | y[1] v[1]; P{TRiP.JF03192}attP2 |
| *Kank* | 33432 | attP2 | y[1] sc[*] v[1] sev[21]; P{TRiP.HMS00319}attP2 |
| *Kirre* | 64918 | attP2 | y[1] sc[*] v[1] sev[21]; P{TRiP.HMC06444}attP2 |
| *Klp3A* | 40944 | attP40 | y[1] v[1]; P{y[+t7.7] v[+t1.8]=TRiP.HMS02192}attP40 |
| *Liprin-α* | 53868 | attP40 | y[1] v[1]; P{TRiP.HMC03183}attP40 |
| *Mad* | 43183 | attP40 | y[1] sc[*] v[1] sev[21]; P{TRiP.GL01527}attP40 |
| *mod(mdg4)* | 32995 | attP2 | y[1] sc[*] v[1] sev[21]; P{TRiP.HMS00849}attP2 |
| *mspo* | 29460 | attP2 | y[1] v[1]; P{TRiP.JF03397}attP2 |
| *MsR1* | 27529 | attP2 | y[1] v[1]; P{TRiP.JF02680}attP2 |
| *nAChRalpha3* | 61225 | attP40 | y[1] v[1]; P{TRiP.JF02750}attP2 |
| *Nadk1b* | 62858 | attP40 | y[1] sc[*] v[1] sev[21]; P{TRiP.HMC05331}attP40 |
| *ND-51L2* | 64536 | attP40 | y[1] sc[*] v[1] sev[21]; P{y[+t7.7] v[+t1.8]=TRiP.HMC05555}attP40 |
| *patched* | 55686 | attP40 | y[1] sc[*] v[1] sev[21]; P{TRiP.HMC03872}attP40 |
| *PhKγ* | 42500 | attP40 | y[1] sc[*] v[1] sev[21]; P{TRiP.GL00214}attP2 |
| *PhKγ* | 35311 | attP2 | y[1] sc[*] v[1] sev[21]; P{TRiP.GL00214}attP2 |
| *Ptp61F* | 56036 | attP40 | y[1] sc[*] v[1] sev[21]; P{TRiP.HMC04344}attP40 |
| *PVRAP* | 61341 | attP40 | y[1] v[1]; P{TRiP.HMJ23233}attP40 |
| *Rabaptin-5* | 52996 | attP40 | y[1] v[1]; P{TRiP.HMJ21709}attP40 |
| *Red herring* | 65085 | attP40 | y[1] sc[*] v[1] sev[21]; P{TRiP.HMC05963}attP40 |
| *Sdc* | 51723 | attP2 | y[1] sc[*] v[1] sev[21]; P{TRiP.HMC03265}attP2 |
| *Sema1a* | 34320 | attP2 | y[1] sc[*] v[1] sev[21]; P{y[+t7.7] v[+t1.8]=TRiP.HMS01307}attP2 |
| *Sema5c* | 29436 | attP2 | y[1] v[1]; P{y[+t7.7] v[+t1.8]=TRiP.JF03372}attP2 |
| *Shark* | 42555 | attP40 | y[1] sc[*] v[1] sev[21]; P{TRiP.HMS04460}attP2 |
| *Shark* | 57017 | attP2 | y[1] sc[*] v[1] sev[21]; P{TRiP.HMS04460}attP2 |
| *sine oculis* | 35028 | attP2 | y[1] sc[*] v[1] sev[21]; P{TRiP.HMS01441}attP2 |
| *Snoo* | 57850 | attP40 | y[1] v[1]; P{TRiP.HMJ21897}attP40 |
| *Sol* | 77432 | attP40 | y[1] sc[*] v[1] sev[21]; P{TRiP.HMC06564}attP40 |
| *Syx5* | 29397 | attP2 | y[1] v[1]; P{TRiP.JF03330}attP2 |
| *Tmtc2* | 44509 | attP2 | y[1] sc[*] v[1] sev[21]; P{TRiP.HMC02899}attP2 |
| *tonalli* | 29372 | attP2 | y[1] v[1]; P{TRiP.JF02536}attP2 |
| *Trpm* | 57871 | attP40 | y[1] v[1]; P{TRiP.JF01236}attP2 |
| *tutl* | 54850 | attP40 | y[1] v[1]; P{TRiP.HMJ21587}attP40 |
| *TwdlW* | 57387 | attP40 | y[1] sc[*] v[1] sev[21]; P{TRiP.HMC04691}attP40 |

| | | | |
|---|---|---|---|
| *Vps-51* | 38280 | attP40 | y[1] sc[*] v[1] sev[21]; P{y[+t7.7] v[+t1.8]=TRiP.HMS01731}attP40 |
